# Spontaneous DNA synapsis by forming noncanonical intermolecular structures

**DOI:** 10.1101/2021.11.11.468201

**Authors:** V.V. Severov, V.B. Tsvetkov, N.A. Barinov, V.V. Babenko, D.V. Klinov, G.E. Pozmogova

## Abstract

We report the spontaneous formation of DNA–DNA junctions in solution in the absence of proteins visualised using atomic force microscopy. The synapsis position fits with potential G-quadruplex (G4) sites. In contrast to the Holliday structure, these conjugates have affinity for G4 antibodies. Molecular modelling was used to elucidate the possible G4/IM-synaptic complex structures. Our results indicate a new role of the intermolecular noncanonical structures in chromatin architecture and genomic rearrangement.

## INTRODUCTION

G-quadruplexes (G4s), noncanonical helical nucleic acid structures composed of stacked G-tetrads, are planar arrangements of four guanine residues bound through Hoogsteen base pairing and stabilised by cations, particularly potassium (1). They adopt several topologies depending on DNA strand orientation (parallel, antiparallel, or hybrid), G-tetrad number, and number of involved strands (intra-vs. intermolecular). Wherever there is a G4-forming sequence in one DNA strand, the complimentary strand always contains (2) a C-rich sequence capable of forming another tetraplex structure, i-motif (IM), which is composed of two parallel-stranded duplexes held together in an antiparallel orientation by intercalated C:CH^+^ base pairs.

Interest in G4s emerges from the growing evidence that they play several important roles in biology. Bioinformatic analyses have shown that putative G4 sites (PQSs) are not randomly distributed along the genome, but cluster at defined regions such as telomeric repeats (3), immunoglobulin switch regions (4), replication origins (5), recombination sites (6, 7), gene promoters (8), and copy number variation breakpoints (9). PQSs are also often present in specific parts of transposable elements (10), particularly in human Alu repeats (11, 12). More than 700,000 PQSs have been found in the human genome (13). This number may be significantly broadened to include imperfect G4s (14) or G4s consisting of guanines from both strands of genomic DNA (15). They are involved in processes such as transcription (16–18), replication (19), recombination (7, 20), and genome instability (21–23). The formation of G4 conformations in the human genome has been visualised *in vivo* (24, 25).

IMs have long been considered a structural curiosity that cannot exist under physiological conditions (IMs are typically stable only at mildly acidic pH due to the requirement for cytosine hemiprotonation, while pH in the nucleus equals ~7.3). However, recent findings indicate that conditions such as negative superhelicity or molecular crowding facilitate IM formation (2). Moreover, sequences with at least five cytosine tracts (thousands of cytosine-rich sequences are present in the human genome) may fold into IM at physiological pH, even in the absence of superhelicity and crowding (26). Finally, *in vivo* IM formation has been visualised in the human nucleus using IM-specific antibodies (27).

Research on biologically relevant G4s has mainly focused on monomolecular structures usually using ssDNAs. Although the G4 and IM structures are well characterised, little is known about their behaviour in dsDNA. Intermolecular noncanonical DNA structures are gaining increasing attention. However, studies in this field are usually performed on model oligonucleotides (28, 29), and investigations of native intermolecular G4s are limited to RNA:DNA hybrids formed during transcription (30). Many G4-related processes, such as recombination, enhancer–promoter interactions (31), chromatin remodelling (32), and chromosomal rearrangements, require a close approach or even physical contact of two DNA chains (or two remote parts of one DNA chain), that is, DNA–DNA synapsis/junction. The participation of various protein factors in these processes is described in detail today, with this passive part being assigned to DNA strands. Simultaneously, the polynucleotide nature allows DNA to form different conformational structures depending on the conditions. We believe that the formation of intermolecular noncanonical structures (G4s and/or IMs) may be an important driver of such processes. However, their folding in dsDNA is not straightforward, so it is difficult to reveal them using methods such as NMR or optical methods. Atomic force microscopy (AFM) (33) is a technique for the direct visualisation of single biological molecules. This method has been used previously to visualise DNA duplexes and triplexes (34), G4s and G-wires (28, 35), IMs (29), and synapsable quadruplex-mediated fibres (36).

In this work, we visualised natural dsDNA fragments containing well-known G4s and model artificial 195 bp DNA duplexes containing (G_3_T)_n_G_3_ sequences (n=1–5) in the middle regions using AFM. AFM scanning of the duplexes revealed intermolecular cruciform and higher-order structure formation that allowed us to assume G4/IM-synaptic complex formation. No signs of such complexes were visible in the AFM images of the control duplexes that lack PQS or its part. The presence of G4 folding in the core of the formed complexes was confirmed by anti-G4-DNA antibody (clone 1H6) (24, 25). Possible nucleotide folding in G4s and IMs, geometry of G4 and IM arrangement relative to each other, as well as the stability of the formed synaptic complexes, were analysed using molecular modelling techniques. Based on the AFM results, we also suggest a mechanism of synaptic complex-promoted DNA strand exchange (recombination).

## MATERIAL AND METHODS

### Synthesis, purification, and MS characterisation of oligonucleotides

Oligonucleotides (ONs) (Table 1) were synthesised using a Biosset ASM-800 DNA synthesiser (Biosset Ltd.; Russia) and standard reagents (Glen Research; USA) following standard phosphoramidite protocols. For synthesising 5′-phosphorylated ONs, solid CPR II (Glen Research) was used. 5′-dimethoxytritylated (DMT) ONs were purified using preparative-scale reverse-phase high-performance liquid chromatography (HPLC) on a 250×4.0 mm Hypersil C18 column (Thermo Fisher Scientific; USA) with detection at λ=260 nm and a linear 7.5–25% acetonitrile gradient in 0.1 M ammonium acetate buffer over 45 min at 50 °C, flow rate: 0.85 mL/min. DMT-protection groups were removed by treatment with 80% acetic acid for 30 min and 5′-phosphorylated ONs after detritylation were treated with 32% ammonium hydroxide for 15 min to eliminate the side chains from 5′-phosphate according to the manufacturer’s instructions. The detritylated ONs were further HPLC-purified in 4– 11.5% acetonitrile gradient in 0.1 M ammonium acetate buffer, ethanol precipitated, and dissolved in 1×TE buffer (10 mM Tris, 1 mM EDTA; pH 8.0) to reach a final concentration of 10 mM. The purity of all ONs was determined to be ≥95% using HPLC. Matrix-assisted laser desorption ionisation time-of-flight (MALDI TOF) mass spectrometry was used to verify the compliance of theoretical and experimental ON masses, as described previously (28). The observed difference between the theoretical and experimental ON masses was less than 3 Da (Table 1).

**Table 1.**
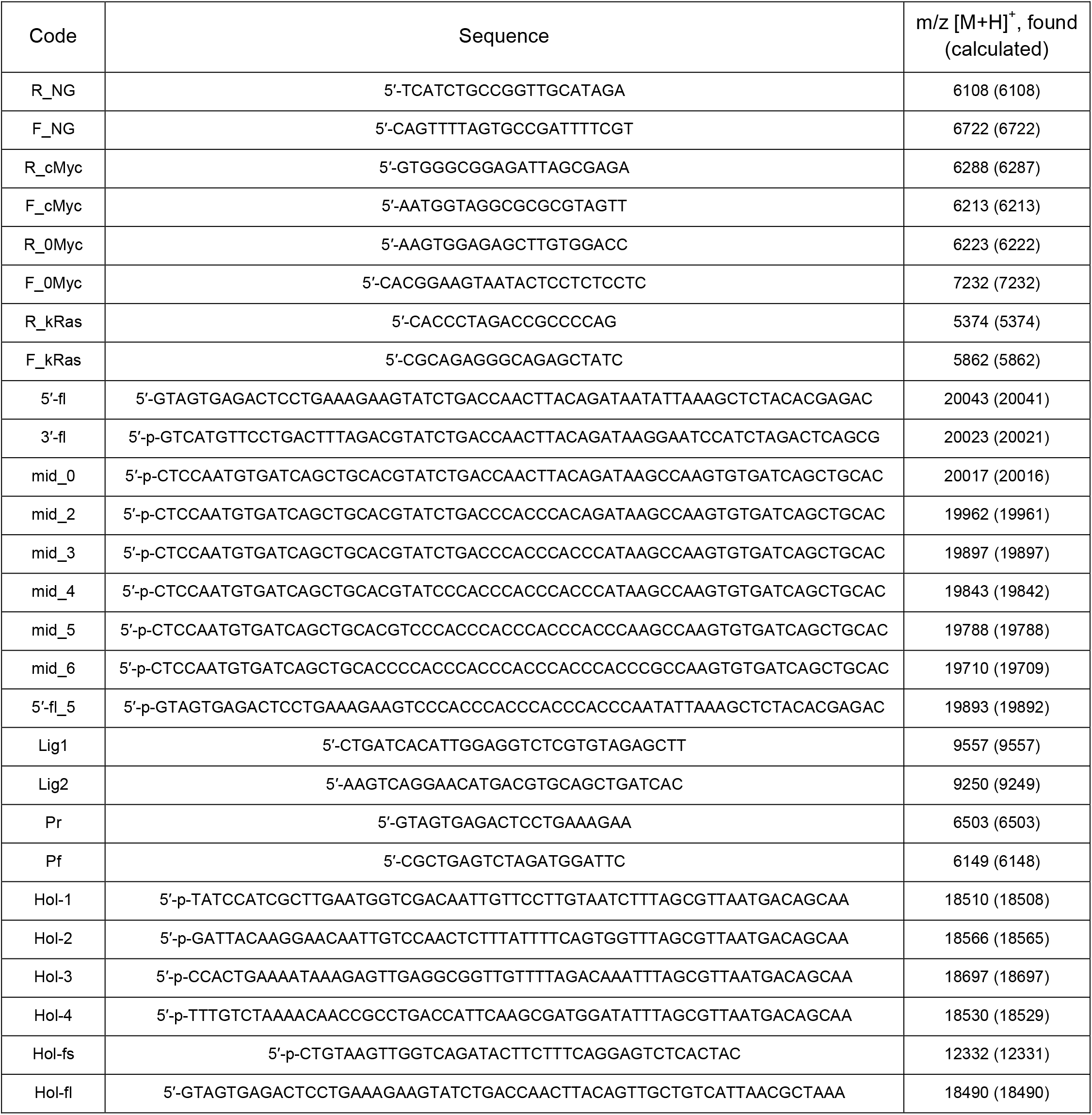
Oligonucleotide sequences and MS data.

### Amplification of human and *N. gonorrhoeae* DNA fragments containing PQS

For amplification of cMyc (201 bp), kRas (208 bp), and control 0Myc (200 bp) duplexes human total genomic DNA (37) was used as template, and *N. gonorrhoeae* (FA1090 strain) total genomic DNA (37) was used as a template for NG duplex (200 bp) production. Amplicons cMyc and NG were amplified using Taq polymerase (Lytech; Russia), and kRas was amplified using the Encyclo GC polymerase kit (Evrogen; Russia), and control duplex 0Myc, with a Screen Mix-HS polymerase kit (Evrogen). Amplifications were performed using a S1000™ thermal cycler (Bio-Rad; USA) under the following conditions: initial denaturation at 97 °C for 3 min, followed by 35 cycles of denaturation at 97 °C for 15 s, annealing at respective temperatures for each primer set (61 °C for R_NG/F_NG and R_kRas/F_kRas, 65 °C for R_cMyc/F_cMyc, and 59 °C for R_0Myc/F_0Myc primer pairs) for 10 s, and elongation at 72 °C for 15 s. The PCR products were separated using electrophoresis on a 2% agarose gel. The amplicons of proper size were excised, gel-purified using the Cleanup Standard kit (Evrogen) according to the manufacturer’s instructions, and washed from the membrane with buffer containing 10 mM Tris-HCl, pH 5.6, and 10 mM KCl (AFM buffer).

### Construction of model DNA duplexes for AFM

DNA duplexes 0, 2m, 3m, 4m, 5m, and 6m (Table S1) were constructed through a two-step PCR (Fig. 2A) using a S1000™ thermal cycler (Bio-Rad). The first PCR amplification was performed in 20 µL reaction mixture, containing 2 µL Lig1, Lig2, Pr, Pf, 5′-fl and 3′-fl, 2 µL 10^x^-dNTPs (Lytech), 2 µL 10^x^-Pfu buffer (100 mM Tris-HCl, pH 8.85, 250 mM KCl, 50 mM (NH_4_)_2_SO_4_, 20 mM MgSO_4_, 1% Tween 20; α-Ferment; Russia) or hand-made buffer containing 200 mM Tris-HCl, pH 8.6; 200 mM LiCl, 25 mM MgCl_2_, 2 µL middle region DNA part (mid_0, mid_2, mid_3, mid_4, mid_5, or mid_6, respectively), 1.5 µL nuclease free water, and 0.5 µL (5 U/µL) Pfu TURBO polymerase (α-Ferment). The reactions were subjected to 20 amplification cycles using the following cycling programme: 94 °C for 10 s, 37 °C for 10 s, and 68 °C for 30 s. The second PCR amplification was performed in 25 µL reaction mixture containing 2.5 µL 10^x^-Pfu buffer or hand-made Li^+^-based buffer, 2.5 µL 10^x^-dNTPs, 0.5 µL first PCR mixture, 1 µL primers Pr and Pf, 17 µL nuclease free water, and 0.5 µL Pfu TURBO polymerase. The reactions were subjected to 30 amplification cycles according to the following cycling programme: 94 °C for 10 s, 50 °C for 15 s, and 68 °C for 30 s; the PCR products were analysed using 10% denaturing (7 M urea) PAGE. The gels were stained with SYBR Gold (Thermo Fisher Scientific) and analysed using a Gel Doc scanner (Bio-Rad). The PCR products were separated using electrophoresis on a 2% agarose gel. The amplicons of proper size were excised, gel-purified using the Cleanup Standard kit (Evrogen) according to the manufacturer’s instructions, and washed from the membrane with AFM buffer (10 mM Tris-HCl, pH 5.6; and 10 mM KCl).

### Sanger sequencing

The amplicon sequences were obtained through Sanger dideoxy sequencing method using Big DyeTM Terminator v.3.1 Cycle Sequencing Kit and ABI Genetic Analyzer 3500XL according to the manufacturer’s instructions (Applied Biosystem; USA).

### The design of an asymmetric Holliday junction

Oligonucleotides Hol-1, Hol-2, Hol-3, and Hol-4 were slowly annealed from 97 °C to 45 °C in 80 µL buffer, containing 20 mM KCl and 10 mM Tris-HCl, pH 7.6 (1.25 pmol/µL each DNA chain). Hol-fl and Hol-fs (5 pmol/µL each DNA chain) solution in 80 µL same buffer was also annealed to 45 °C. Subsequently, these two solutions were quickly mixed and slowly annealed to room temperature (≈25 °C). The folded structure was ligated using T4 DNA ligase (Thermo Fisher Scientific) according to the manufacturer’s instructions. The Holliday junction was separated analogically to the PCR products and dissolved in AFM buffer or AFM buffer supplemented with 2 mM MgCl_2_.

### AFM sample preparation, image acquisition and processing

AFM were performed on freshly cleaved graphite surfaces rendered hydrophilic with an amphiphilic modificator (CH_2_)_*n*_(NCH_2_CO)_*m*_-NH_2_ (34). DNA samples were diluted 20–40 times with AFM buffer, applied on the substrate surface, incubated for 5–15 s, and removed with a nitrogen stream, thus drying the surface for imaging in air. The low salt concentration of the dilution buffer allowed us to eliminate a rinsing step from the sample preparation procedure, which could otherwise alter the folding of synaptic structures. All experiments were performed at least in triplicate. AFM imaging was performed using a multimode AFM instrument with an NTEGRA Prima controller (NT-MDT; Russia) in tapping mode with 1 Hz scan rate and a typical free amplitude of several nanometres. All measurements were performed in air using supersharp cantilevers grown on the tips of commercially available standard silicon cantilevers using a chemical vapour deposition process (spike diameter: approximately 1 nm) (34). FemtoScan Online software (ATC; Russia; http://www.femtoscanonline.com) was used to filter and present the AFM data. Standard algorithms for AFM image flattening were used (subtracting the quadric surface and averaging by lines), and no algorithms for resolution improvement were used; thus, the raw AFM images are presented in this study. SPM Image Magic software (https://sites.google.com/site/spmimagemagic, Alex Kryzhanovsky) was used to semi-automatically analyse the ON heights. The analysis consisted of two steps: the individual particles were identified automatically on the images by the local maxima, and their heights were calculated with respect to the local background surrounding the particles. The results of the automatic analysis were filtered manually when necessary.

### Molecular modelling and molecular dynamic simulation

All 3D models of the studied structures were built using the molecular graphics software package Sybyl-X software (Certara; USA) using the following strategy. Initially, models of the required duplexes, quadruplexes, and IMs were created. Further, the created models were located relative to each other in the required geometry and connected. At each stage, molecular mechanical optimisation was performed to eliminate the van der Waals overlap, which could occur during a certain step. The molecular mechanical optimisations were performed using Sybyl-X and Powell’s method with the following settings: parameters for intermolecular interactions and the values of partial charges: taken from force field amber7ff99, non-bonded cut-off distance: 8 Ǻ, effect of the medium: dielectric constant of 4, the number of iterations: 1000, simplex method for initial optimisation, and 0.05 kcal*mol^−1^*Å^−1^ energy gradient convergence criterion. The stability of the created models was tested by molecular dynamics using Amber 20 software (38). The MD simulations in the production phase were performed using constant temperature (T=300 K) and pressure (p=1 atm) over 4 ns. To control the temperature, a Langevin thermostat was used with 1 ps^−1^ collision frequency. Influence of the solvent simulated with the application model of water molecules OPC3 (39). K^+^ ions were used to neutralise the negative charge of the DNA backbone. The parameters needed for the interatomic energy calculation were taken from the force fields OL15 (40, 41).

## RESULTS

### G4/IM-synaptic structure formation by DNA duplexes containing PQS

We studied DNA duplex fragments of human and *N. gonorrhoeae* genomes (≈200 bp) with PQS in its middle regions using high-resolution AFM. DNA samples containing well-known G4s of two oncogene promoters: cMyc duplex (201 bp) including Pu27 PQS of cMyc promoter NHE III_1_ element (42), kRas duplex (208 bp) including PQS in GA-element of KRAS gene promoter (43), and G4-forming sequence located upstream of the *N. gonorrhoeae* pilin expression locus (NG duplex, 200 bp) required for pilin antigenic variation (7), were chosen. 0Myc (200 bp) sequence, located near the cMyc fragment of the human genome and without PQS, was used as a control. The sequences are listed in Table S1.

The DNA samples were obtained through PCR amplification of human or *N. gonorrhoeae* (FA1090 strain) total genomic DNA (37). The primers (Table 1) were selected using the Primer-BLAST tool (https://www.ncbi.nlm.nih.gov/tools/primer-blast/). The amplicons were separated by agarose gel electrophoresis and dissolved in a buffer containing 10 mM Tris-HCl (pH 5.6) and 10 mM KCl (AFM buffer).

An AFM image of the control duplex 0Myc is shown in Fig. 1A. As seen in the figure, only separate DNA molecules (height 1.0±0.1 nm and length 65±3 nm) are found, which corresponds with DNA length in solution (44). Some of the molecules have melted areas, such as single-stranded loops and tails. This is the main difference between the molecules.

**Fig. 1.**
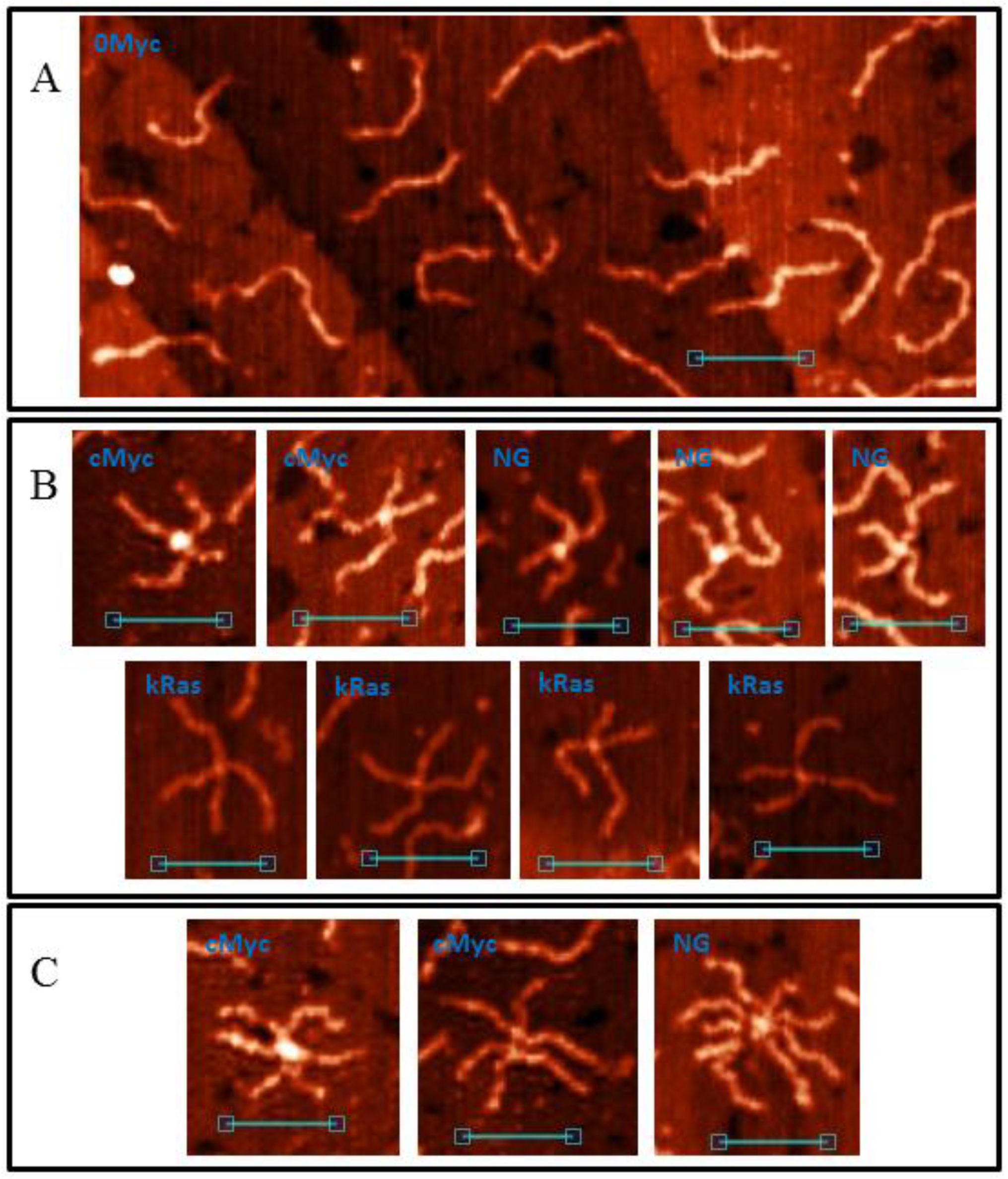
AFM images of natural dsDNA fragments. **A**. AFM image of 0Myc duplex; **B**. Cruciform structures, formed by PQS-containing duplexes; **C**. Complex G4/IM-synaptic complexes formed by more than two molecules. Scale bars: 50 nm.

DNA samples cMyc, kRas and NG, containing PQSs in their middle regions, are also presented mostly by separate molecules, but with this the images contained cruciform (Fig. 1B) and higher-order structures (Fig. 1C), we called them G4/IM-synaptic complexes. The measured heights of the central cores varied from molecule to molecule and were from practically absence of elevation in comparison to the duplex arms up to 2.5 nm. Thus, the two DNA duplexes may form different structure junctions. In addition, some complexes are joined not by their middle regions, which are the PQSs. These sequences are GC-rich and contain areas where two or more G_3_-tracks are divided by less than seven nucleotides. They may also form synaptic complexes through intermolecular G4 or IM.

To investigate G4/IM-synaptic complex structures in detail, we synthesised a model set of 195 bp DNA duplexes containing varying numbers of G_3_-tracks ((G_3_T)_n_G_3_, n=1–5) within statistical duplex media. They were visualised by AFM, and the images were compared with the results of possible synaptic complex structures developed using molecular modelling techniques.

### Synthesis of DNA constructs for AFM

The sequences of 0, 2m, 3m, 4m, 5m, and 6m dsDNA-constructs are given in Table S1. They differ only in their middle section, so they were obtained using a universal two-step PCR (Fig. 2A, oligonucleotides used for amplification are shown in Table 1). The analytical PAGE of this two-step synthesis is shown in Figure 2B. Lines 3, 5, 7, 9, 11, and 13 show the results of PCR conducted using commercial buffer (100 mM Tris-HCl, pH 8.85, 250 mM KCl, 50 mM (NH_4_)_2_SO_4_, 20 mM MgSO_4_, and 1% Tween 20). PCR amplification of 0, 2m, and 3m was efficient under such conditions, but the efficiency decreased significantly when amplicons contained sequences capable of intramolecular G4 folding (4m, 5m, and particularly 6m); moreover, we observed a discrete ~100 bp product formation along with the expected 195 bp amplicon (lines 9, 11, 13). A similar appearance of short PCR products was observed during amplifying the *vlsE* region from *B. burgdorferi* (20), and it is common knowledge that the sequences containing PQS are difficult PCR templates. Amplification of such sequences (particularly construction of PQS-containing duplexes from oligonucleotides using PCR) often requires intensive optimisation and/or the use of PCR additives (45). PCR buffer conditions (pH>8) generally prevent IM formation, but the PCR buffer usually contains K^+^ and NH_4_^+^ salts, which facilitate G4 folding. G4s are reportedly unstable in Li^+^ salts (46); therefore, we substituted all K^+^ and NH_4_^+^ salts with LiCl and performed PCR amplification in a buffer with 20 mM Tris-HCl, pH 8.6, 20 mM LiCl and 2.5 mM MgCl_2_. Lines 2, 4, and 6 (Fig. 2B) in comparison with lines 3, 5, and 7 show that the amplification efficiency in Li-based buffer is generally lower than that in commercial buffer, but the amplification of sequences containing intramolecular G4s (lines 8, 10, and 12) was more efficient (lines 9, 11, 13) and similar to that of sequences not containing PQSs (lines 2, 4, and 6). This similarity may be used to address the problems associated with allele dropout during PCR of single nucleotide polymorphisms containing PQS (45). After amplification, duplexes 0, 2m, 3m, 4m, 5m, and 6m were separated using agarose gel electrophoresis and dissolved in AFM buffer. Similarly, duplex 5l with PQS not in the middle, but shifted, was obtained: 5-fl_5, mid_0, and 3′-fl were used as the main building blocks in the first PCR step.

**Fig. 2.**
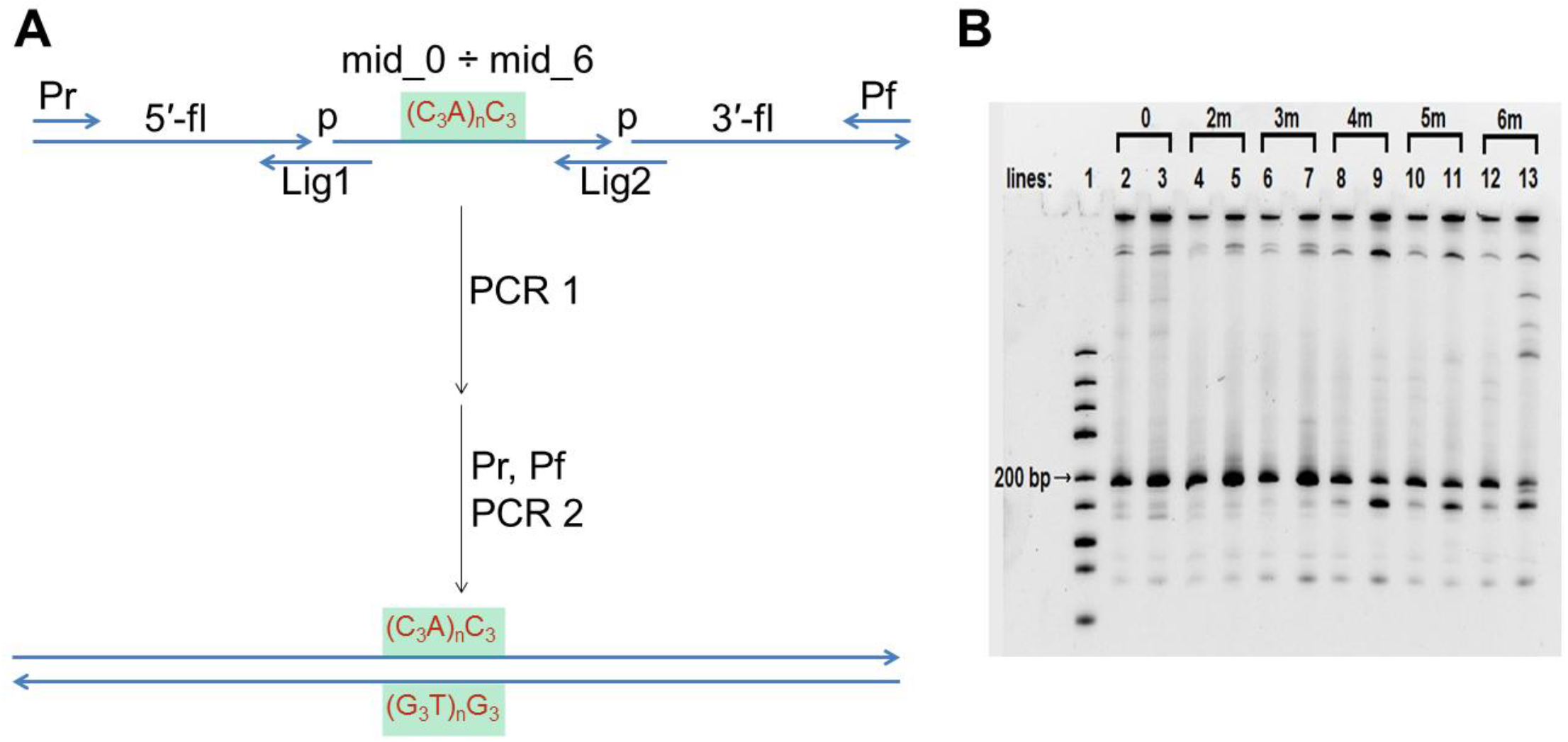
Synthesis of model DNA duplexes. **A**. Scheme of two-step PCR synthesis of model DNA constructs; **B**. Analytical PAGE of amplification. Lines 3, 5, 7, 9, 11 and 13 – PCR in the commercial buffer; lines 2, 4, 6, 8, 10 and 12 – PCR in the in-house Li-based buffer.

### Structure of G4/IM-synaptic complexes

DNA duplex 0 without G_3_-tracts in its sequence was used as a control. Its AFM images are similar to those of 0Myc duplexes, the control images indicate the presence of 62±3 nm long separate DNA molecules, which may contain melted areas.

DNA duplexes 2m and 3m may form synaptic complexes with two the simplest structures, where the duplexes join through intermolecular G4 or IM formation. Molecular modelling of the first case (joining through G4) is shown in Fig. 3A (hereafter, noncanonical shapes are highlighted with wider rendering; K^+^ ions, which stabilise G4s, are highlighted in magenta). Such structures must have a stiff G4 core and duplex arms must disperse symmetrically. Most of the synaptic complexes, visualised by AFM, are of this geometry (Fig. 3B), with a 1.1–1.4 nm core structure height. Angles between duplex arms vary and depend on the manner in which the molecular associate lies on the substrate surface. The morphology of a stable synaptic complex with joining through IM differs from that discussed above as the folded IM serves as a dividing bridge between two duplexes (Fig. 3C). AFM images of these molecular associates are shown in Fig. 3D. The core part, dividing the duplexes, is 5–7 nm long and has 1.2±0.1 nm height. Molecular modelling also revealed another possible stable IM structure in duplex media (Fig. 3E). The formation of such a structure is impossible is this study because of the need for interlacing of DNA chains, but theoretically may fold while the Holliday structure moving, thus influencing the recombination processes. The difference between 2m and 3m in the formation of such simple two-duplex complexes is that both available G_3_ (or C_3_) blocks in the 2m sample participate in synaptic structure formation; however, different two of the three blocks in the 3m sample may form the same synaptic structure, and the resolution of AFM is not sufficient to discriminate between them.

**Fig. 3.**
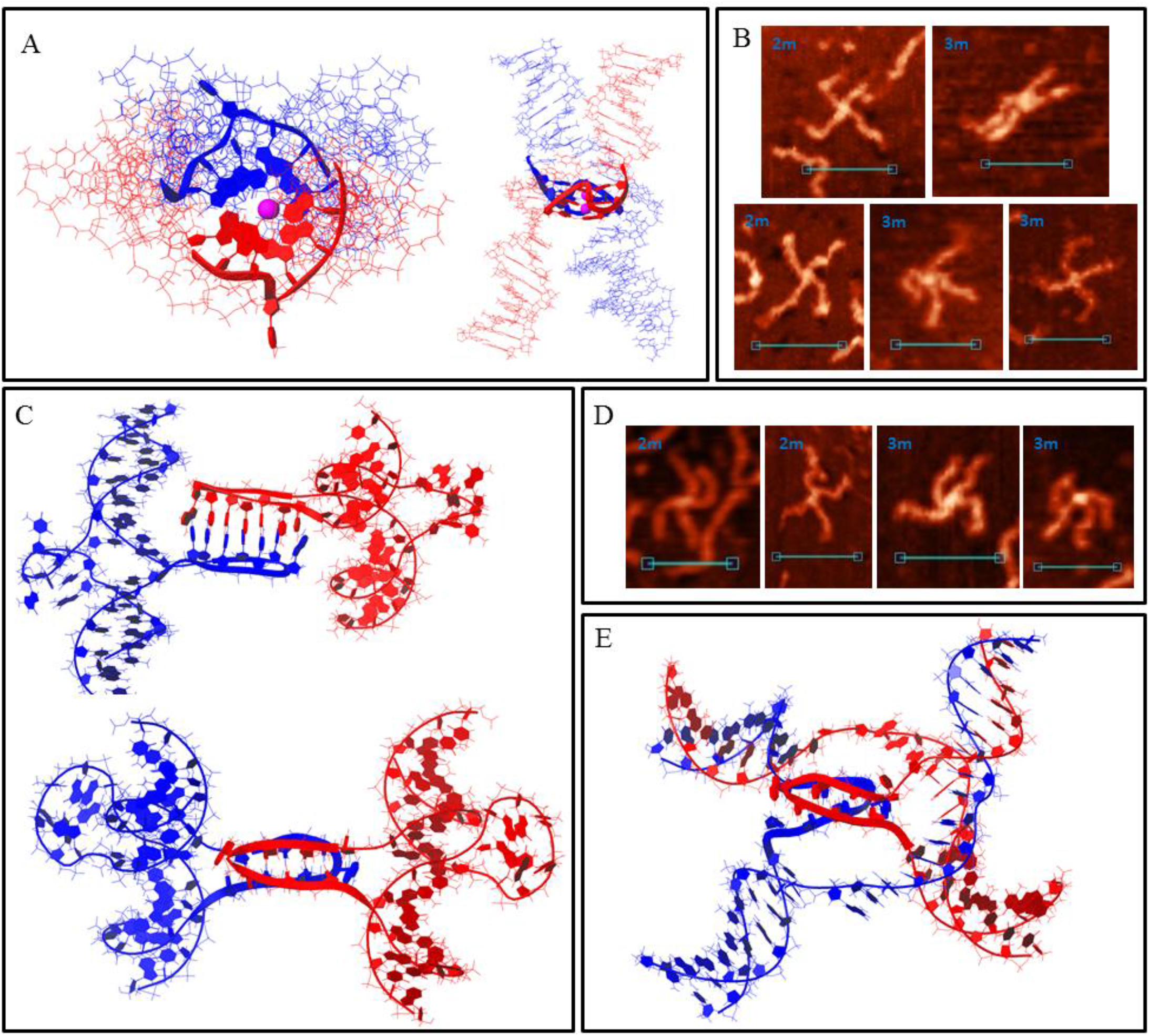
Bimolecular G4/IM-synaptic complexes formed by 2m and 3m samples. **A**. Molecular model of duplex joining through intermolecular G4 folding. **B**. Examples of the complexes consistent with model **A**, revealed by AFM. Scale bars: 50 nm. **C**. Molecular model of the joining through intermolecular IM folding. **D**. Examples of the complexes consistent with model **C**, revealed by AFM. Scale bars: 50 nm. **E**. Molecular model of possible intermolecular IM formation upon Holliday structure moving.

When one G-rich (or C-rich) chain of a DNA duplex participates in a synaptic complex formation, the other free C-rich (or G-rich) chain may interact with the third duplex molecule, thus forming a multimeric G4/IM-synaptic complex. Such structures, formed of three duplexes, have been found in the 2m and 3m samples (Fig. 4A). These associates have clearly distinguishable gaps within duplex the junctions and are not symmetrical as one core part is more stiff and high, while the other is lengthy. A stable structure that fully corresponded to the AFM images is shown in Figure 4B. Molecular modelling explains the gap between the two folds and predicts that G4 is located perpendicular to the IM. Theoretically, the joining of next molecules and synaptic complex growth may go further by the described method (for example, the tetramolecular junction is shown in Figure S1A), but we did not find such large complexes for the 2m sample. This may be related to the insufficient stability of such large complexes under AFM conditions. However, multimeric structures formed of more than three duplexes were revealed for the 3m sample. Moreover, they are formed even more often than the smaller complexes described above. Molecular modelling predicts another possibility of joining, where it forms a G4 chain with no need for IM folding. When the synaptic complex is composed of three duplexes, the synaptic core contains two G4s, the first of which consists of three G_3_ blocks of the first duplex molecule and one of the second, the remaining two G_3_ blocks of the second duplex form G4 with two G_3_ blocks of the third duplex (Fig. S1B). The G4s in this case are located in close proximity and perpendicular to each other. Trimolecular associates of such geometry were not revealed by AFM, but tetramolecular complexes folded according to the same logic of G4 chain formation (Fig. 4C) were found (Fig. 4D). The central G4 is formed by two G_3_ blocks from each central duplex. The remaining third blocks formed side G4s with three G_3_ blocks of side duplexes. Eight-arm synaptic complexes have 1–3 G4 cores depending on the way the molecular associate lies on the substrate surface. At the front view of the associate (as shown in Fig. 4C), all three G4s are distinguishable and have height of 1.4–1.6 nm (bottom panel in Fig. 4D). The green arrow indicates the viewpoint from which the complex appears as in the middle AFM scan with core heights 1.6–2 nm. The red arrow indicates the viewpoint where the three G4s lie on each other (upper AFM scan, core height 2.8 nm). Further synaptic complex growth by this scheme is impossible, but there is no steric hindrance for free C-rich chains of side duplexes to form IM with the next duplexes (Fig. 4E), thus forming stable 5- or even 6-duplex complexes. Most of these large complexes were disrupted during absorption to the substrate surface or by cantilever while scanning and observed as a shapeless mixture of synaptic complexes (Fig. 4F), but one good example, confirming this possibility, was found (Fig. 4F, bottom panel).

**Fig. 4.**
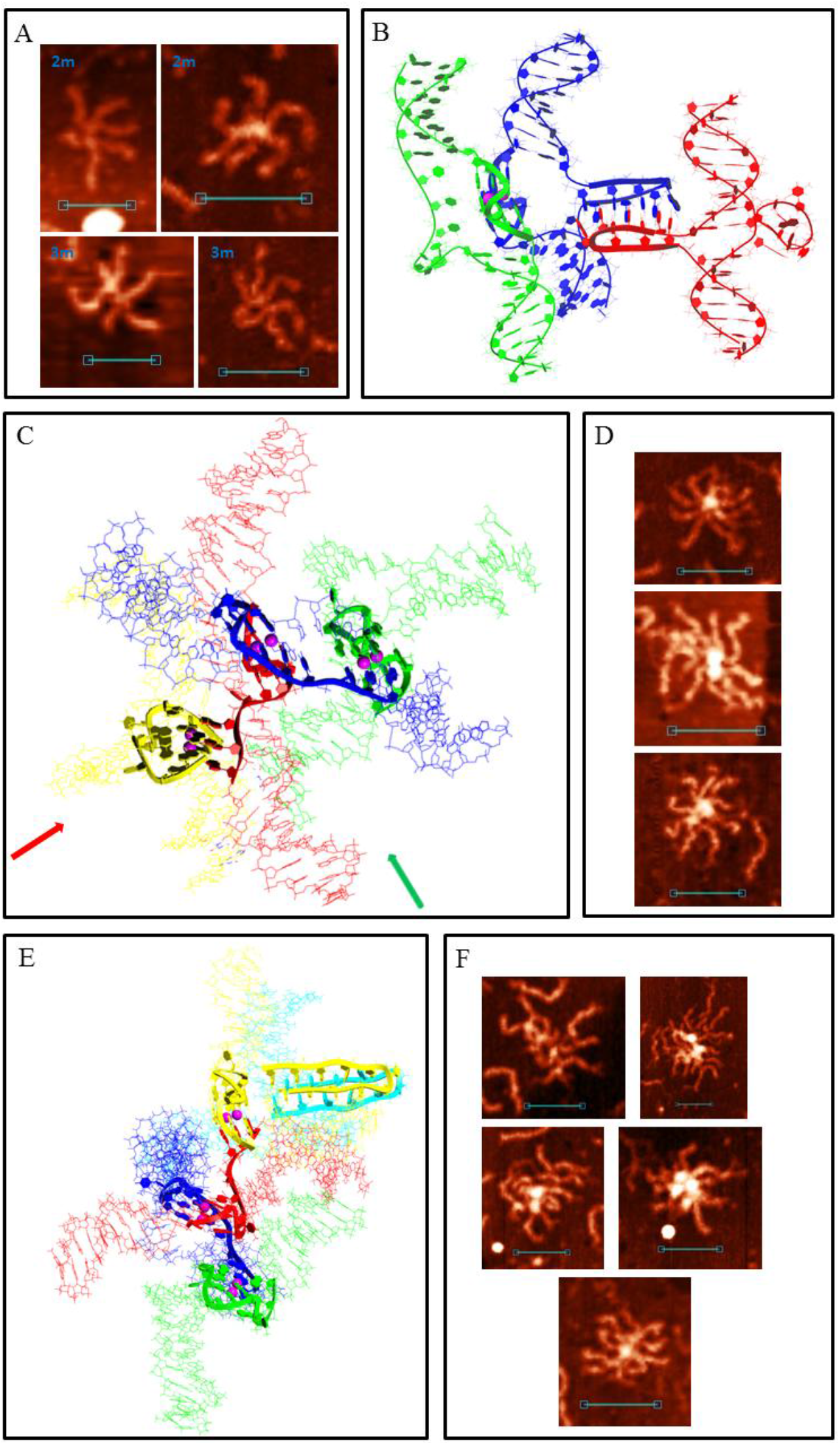
Multimeric G4/IM-synaptic complexes formed by 2m and 3m samples. **A**. Examples of trimolecular complexes formed through intermolecular G4 and IM folding. Scale bars: 50 nm. **B**. Molecular model of the structure, consistent with **A. C**. Molecular model of the tetrameric complex formed by 3m sample through the formation of three intermolecular G4s. The red and green arrows show the viewpoints from which the complexes are visible in the following examples. **D**. Examples of the tetrameric complexes, revealed by AFM. Scale bars: 50 nm. **E**. Molecular model of the synaptic complex formed by additional duplex joining with the previous structure through intermolecular IM formation. **F**. Examples of the complexes, consistent with model **E**, revealed by AFM. Bottom image represents the only example of the complex that remained intact (was not disrupted) during the AFM experiment. Scale bars: 50 nm.

Sample 4m may form all synaptic complexes formed by 2m and 3m duplexes, and many such complex types (as bimolecular and multimeric) with the same parameters (core length and height), have been observed using AFM (Fig. S2). With this by the same scheme/logic of bimolecular synaptic complex formation via intermolecular G4 or IM folding, 4m duplexes may join through more complex structures where all four G_3_ or C_3_ blocks participate. Molecular modelling of the association with the help of G-rich chains is shown in Figure 5A. Joining involves the formation of an interlocked G4-dimer, whose structures at the single-chain level are thoroughly described previously (28). Residual single-stranded C-rich chains wrap the G4-core. In the AFM images (Fig. 5B), such complexes are similar to those described above for joining through G4 (compare with Fig. 3B); but in this case, the core is larger and has 1.6±0.2 nm height. Similarly, instead of intermolecular IM formed by 2m and 3m samples (Fig. 3C), the presence of the fourth CCC-block leads to folding of two IMs, divided by bulged thymidine residues between them. Molecular modelling (Fig. 5C) predicts that thymidines do not fold into a tetrad or any other structured form. It also predicts that the released G-rich chains are able to fold into intramolecular G4s opposite intermolecular IMs. AFM images of synaptic complexes referring to this structure are shown in Figure 5D. The core part has 10±2 nm length and 1.2±0.1 nm height. It is unclear whether G4 folded in most scans, but in the fourth AFM image (from left to right) the molecular associate fell into a surface gap, which corresponds to its core size. This makes border structures distinguishable from the central 2IMs part. It is clearly visible that one G-rich chain forms no secondary structure (red arrow) and the second forms discernible globule (G4, green arrow). Therefore, we concluded that G4 formation may occur in this synaptic complex type. Another feature of the structure is the twist of the 2IM bridge, which is not notable in AFM images because synaptic complexes tend to press down to the substrate surface during the AFM experiment, but we found one clearly visible twist (fifth image).

**Fig. 5.**
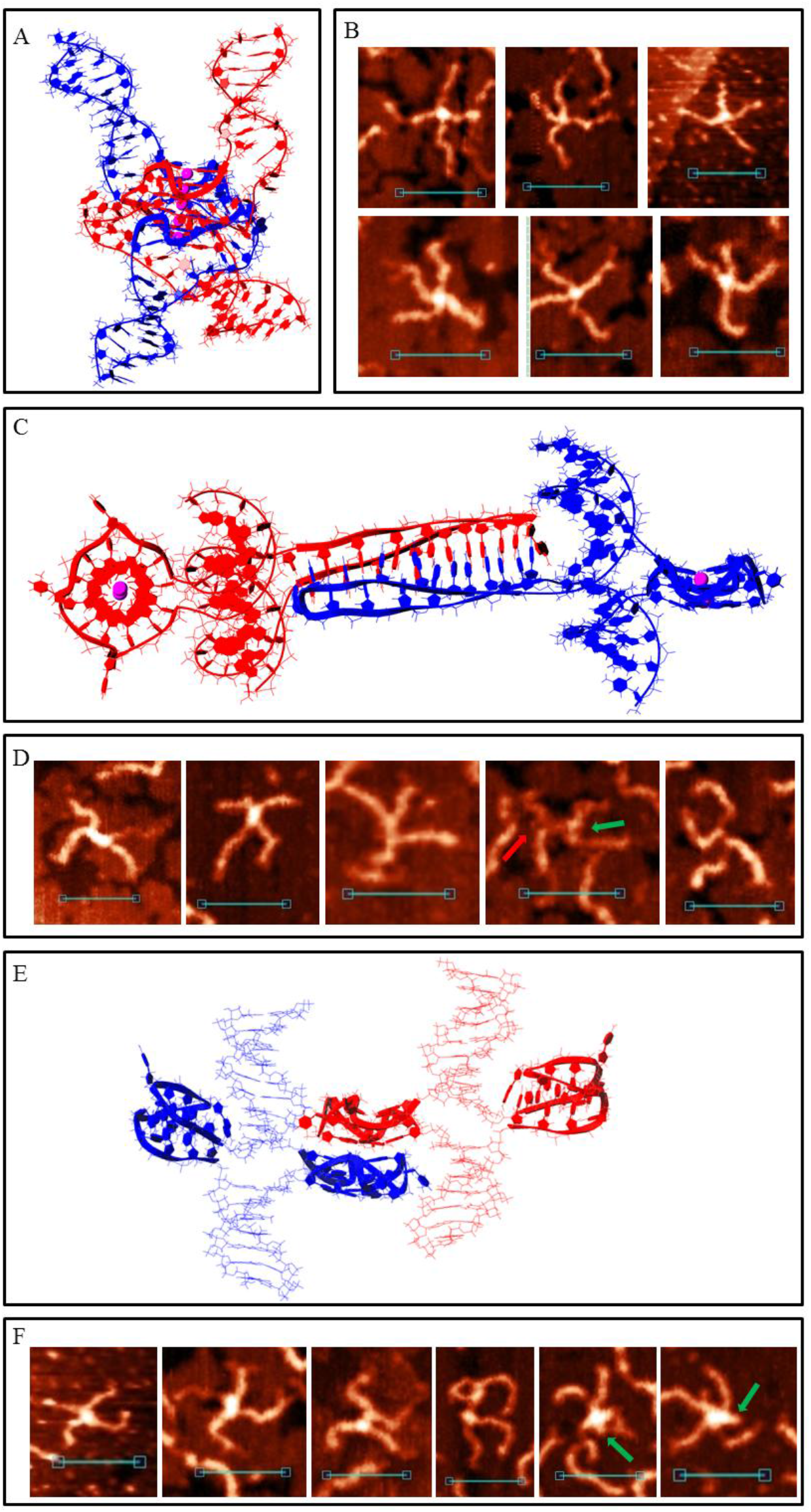
Bimolecular G4/IM-synaptic complexes formed by 4m sample. **A**. Molecular model of duplex joining through interlocked G4-dimer folding. **B**. Examples of the complexes, consistent with model **A**, revealed by AFM. Scale bars: 50 nm. **C**. Molecular model of duplex joining through two intermolecular IMs. **D**. Examples of the complexes, consistent with model **B**, revealed by AFM. Red arrow indicates the free G-rich chain, and green arrow indicates the folded G4 structure. Scale bars: 50 nm. **E**. Molecular model of duplex joining through stacking of intramolecular G4s. **F**. Examples of the complexes, consistent with model **E**, revealed by AFM. Green arrows indicate folded IM structures. Scale bars: 50 nm.

The possibility of intramolecular G4 or IM formation expands the diversity of possible synaptic complex structures. This opens the way for another assembly type through stacking of intramolecular G4s. For such complexes of such type, molecular modelling predicts that the released C-rich chains are able to fold into IMs (Fig. 5E). Previously, it has been shown that G4 and IM formation is mutually exclusive in duplex media (47); however, this has not been studied in detail. In this work, the PQS sequence is another, and theoretically, their simultaneous formation is possible, at least in synaptic complex structure. The most stable stacking conformation occurs when the duplexes are the most distant, so they (and possible IMs) are opposite to each other relative to the G4–G4 core. AFM images of such assemblies must be (in case of IMs are not folded) similar to the previously described case of joining through intermolecular G4 (Fig. 3B). We recognise them because G4 stacks are less stable than G4 structures and tend to disrupt at the substrate surface. Consequently, we observed different stages of stack decay (Fig. 5F). The height of the G4 cores (irrespective of decay or stacking level) is 1.2–1.4 nm. When the complex was not disturbed (fifth and sixth AFM images), at least one of the two IMs fold (green arrows). The height of the central cores in this case is 1.4–1.6 nm, while that of the side IMs is ≈1.2 nm. It should be noted that no IM was observed opposite to the folded G4 in structures with disrupted stacking.

The diversity of multimeric synaptic complex structures grows too in the case of a 4m duplex. Of those observed for the 2m and 3m samples, only the trimolecular associates formed by G4 and IM folding, were found here (Fig. S2). There were also centrally symmetric assemblies with 7–8 duplex arms and 1.8–2.2 nm core height (Fig. 6A). We ascribed them to the tetrameric complex formed completely due to intramolecular G4 stacking (Fig. 6B). This structure explains the absence of the eighth arms in most cases as they lie down and are hidden from the cantilever during AFM. For the same reason, only the upper IM was visible (green arrow). Trimolecular assemblies formed through stacking were not revealed by AFM and molecular modelling confirmed that such associates are not stable. Therefore, we concluded that tetramolecular complexes are not built by the consecutive joining of duplexes, but may form only by stacking two dimers. In this case, the central stacking goes perpendicularly (view from the top is symmetrical). Another peculiar tetrameric complex is shown in Figure 6C. We did not make a model of its structure, but it is clearly visible that it was folded by a combination of G4–G4 stacking and 2IMs (as in Fig. 5C). Therefore, different methods of joining lead to the formation of large multimeric complexes with indistinguishable structures (Fig. 6D).

**Fig. 6.**
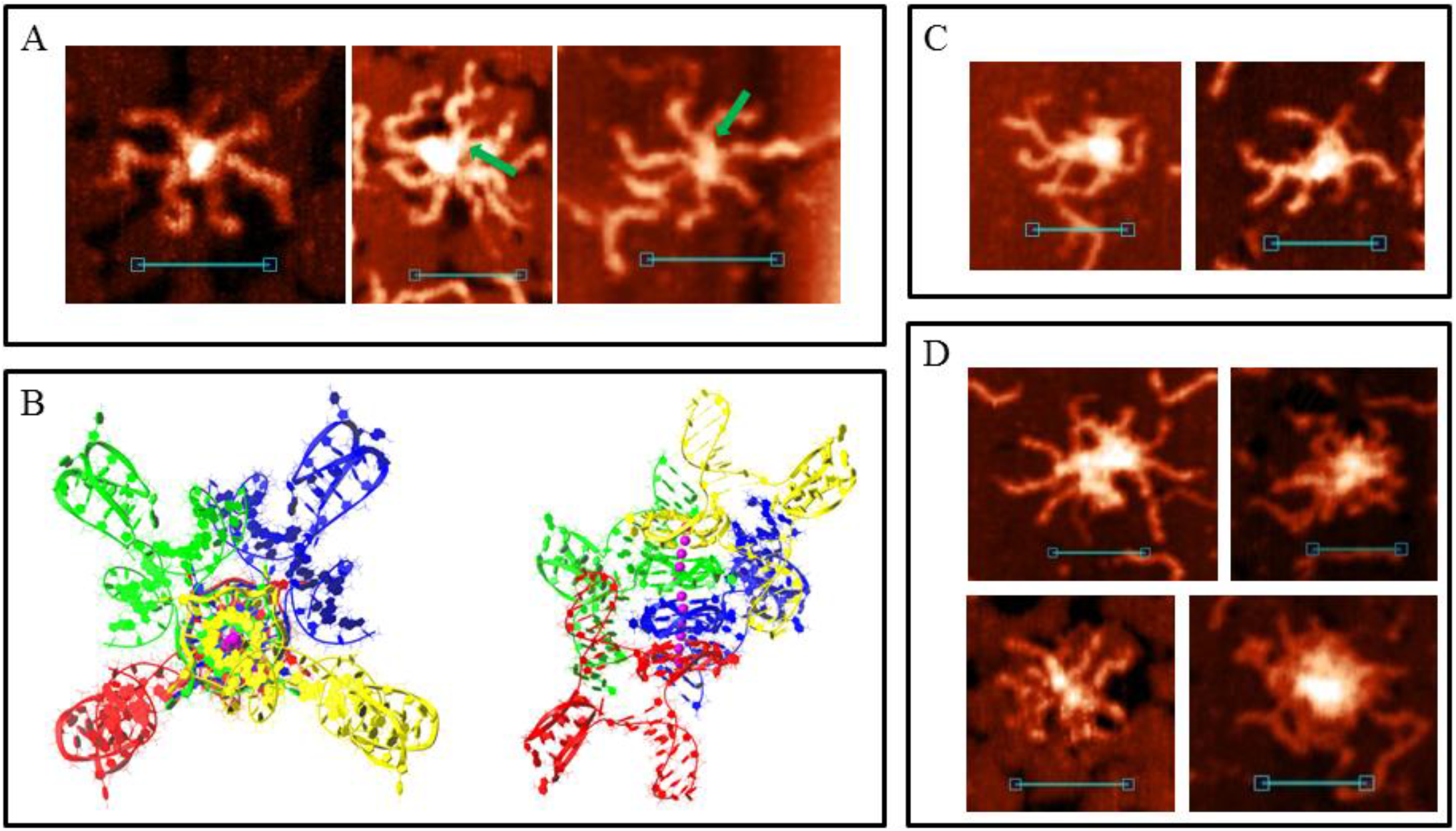
Multimeric synaptic complexes formed by 4m sample. **A**. Examples of the complexes, formed through stacking of four intramolecular G4s, revealed by AFM. Green arrows indicate folded top IM structures. Scale bars: 50 nm. **B**. Molecular model of the structure, consistent with **A** (top and side views). **C**. Tetramolecular synaptic complexes formed by 4m sample through a combination of G4-G4 stacking and the formation of two intermolecular IMs separated by thymidine residues. **D**. Big complexes with indistinguishable structures. Scale bars: 50 nm.

Most common synaptic complex structures formed by the 5m and 6m samples were identical to those formed by the 4m sample (Fig. S3A, Fig. 5). Complexes with structures depicted in Fig. 3 are twice as rare. The AFM images of these complexes may differ from that of those formed by 2m, 3m and 4m samples only because different G_3_/C_3_-blocks may fold in these cases; therefore, duplex arm lengths will differ (particularly for complexes with structures of 2m and 3m forms, Fig.S3B). The presence of an additional G_3_/C_3_-block may transform one or two (in the case of a 6m sample) middle blocks to a 5-base (TG_3_T/AC_3_A) or a 9-base loop. Such complexes based on intermolecular IMs (Fig. 5C) are not distinguishable from the initial ones (Fig. 5D) in AFM images or absent, so we did not develop their molecular models. Complexes with one 5-base loop based on G4-G4 stacking (Fig. 5E) in the AFM images were attributed to those shown in Figure 7A. Molecular modelling predicts that any inner G_3_-block may become a loop, but at the most stable variant (shown in Fig. 7B) at one stacked G4 the second block (from the 5′-end) is looped out, and at the other G4 – the fourth. Both IMs are folded in such a way that third C_3_-blocks are looped out. These complexes are more stable than the initial ones shown in Fig. 5, AFM also confirms that: there are not so many pre-folded (or partly destroyed) examples, and one or both IMs are usually present. The quantity of folded IMs is distinguished by the core geometry (stretched in case of one IM and bent in case of two); single-stranded not IM-folded loops are usually undetectable and only one example was found (right image in Fig. 7A). The height of the core part is 1.2–1.4 nm. Molecular modelling indicates that for structures of this type with two long loops in each chain (6m sample) at the G4–G4 core (as in the case of one 9-base loop), IM folding by central C_3_ blocks (side blocks serve as stems between IMs and other parts of the complex) without long loops is most energetically favourable (Fig. S4). The AFM images of such complexes must be similar (may be not distinguishable) to that of those with one 5-base loop (Fig. 7A), may be a bit wider, but we did not find a wider 6m sample structure with the same geometry, so it cannot be said exactly that they fold under the AFM conditions. Duplexes with at least one additional G_3_/C_3_-blocks may also join through interlocked G4-dimer (Fig. 5A), but the presence of 5-base loops and longer C-rich single chains that wrap G4-dimer (Fig. 7C) widens the core in AFM images (2–2.5 nm height, Fig. 7D). Synaptic complexes of such structures are quite abundant in the 5m and 6m samples. Analogically, the 6m sample is theoretically capable of forming structures with two 5-base loops at each G4 (or one 11-base loop), but AFM shows no synaptic complexes with the same geometry, but with a wider and higher (>2.5 nm) core part. It should be noted that there is also the possibility of folding several complexes with mixed topology, where, for example, one duplex forms a 5-base loop, but the second uses only neighbouring blocks in synaptic complex formation. We did not model all these possibilities, partly due to their multiplicity, but tried to consider the basic principles of their composition. For example, we may suggest that the large height variation of complexes, depicted in Fig. 7D (2–2.5 nm), is connected with the formation of such mixed structures.

**Fig. 7.**
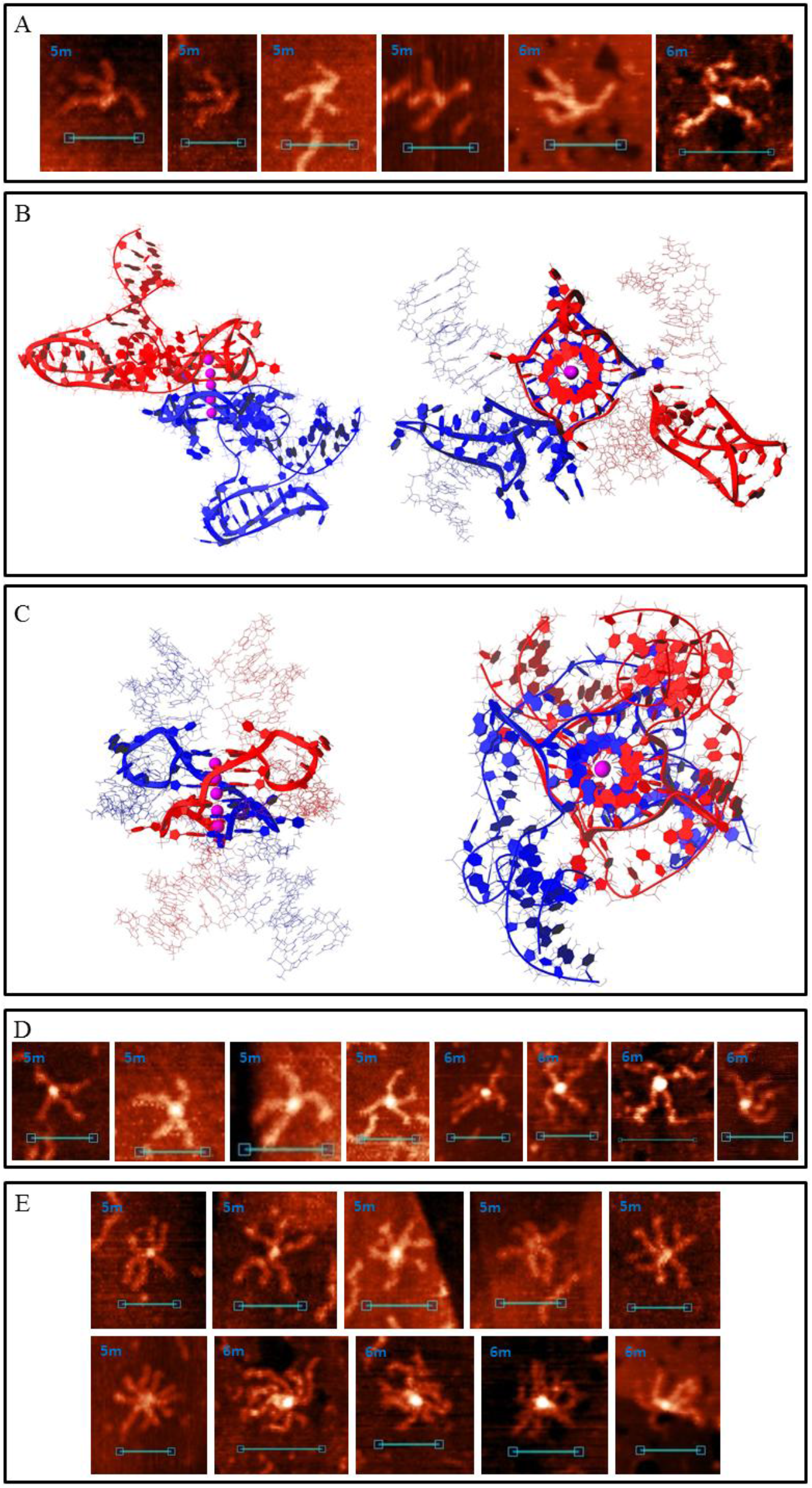
G4/IM-synaptic complexes formed by 5m and 6m samples. **A**. Examples of bimolecular complexes, formed through stacking of intramolecular G4s, which contain 5-base loops, revealed by AFM. Scale bars: 50 nm. **B**. Molecular model of the structure, consistent with **A** (with two IMs folded). **C**. Molecular model of duplex joining through the interlocked G4-dimer, in which each G4 contains 5-base loops. **D**. Examples of the complexes, consistent with model **C**, revealed by AFM. Scale bars: 50 nm. **E**. Examples of small multimeric G4/IM-synaptic complexes, formed by 5m and 6m samples.

Multimeric synaptic complexes, formed by 5m and particularly 6m sample, tend to grow and usually include dozens of duplexes. Even for relatively small complexes, including 3–5 molecules, each sample has its own structure, formed by mixing different joining methods (Fig. 7E).

Moreover, 5s duplex (Table S1) was constructed using the same universal two-step PCR scheme (Fig. 2). Five G_3_/C_3_-blocks in it are not located in the middle of the duplex molecule, but shifted. AFM images indicate synaptic complexes formed by the 5s sample (Fig. 8). It is clearly visible that the joining crosshair is also shifted, as for bimolecular and multimeric complexes. The duplex arm length ratio is about 4–7:1 (depending on which G_3_/C_3_-blocks participate in complex folding), which is consistent with the PQS position in the DNA sequence. This is an additional proof that duplexes are joined by G4/IM-synaptic complex folding.

**Fig. 8.**
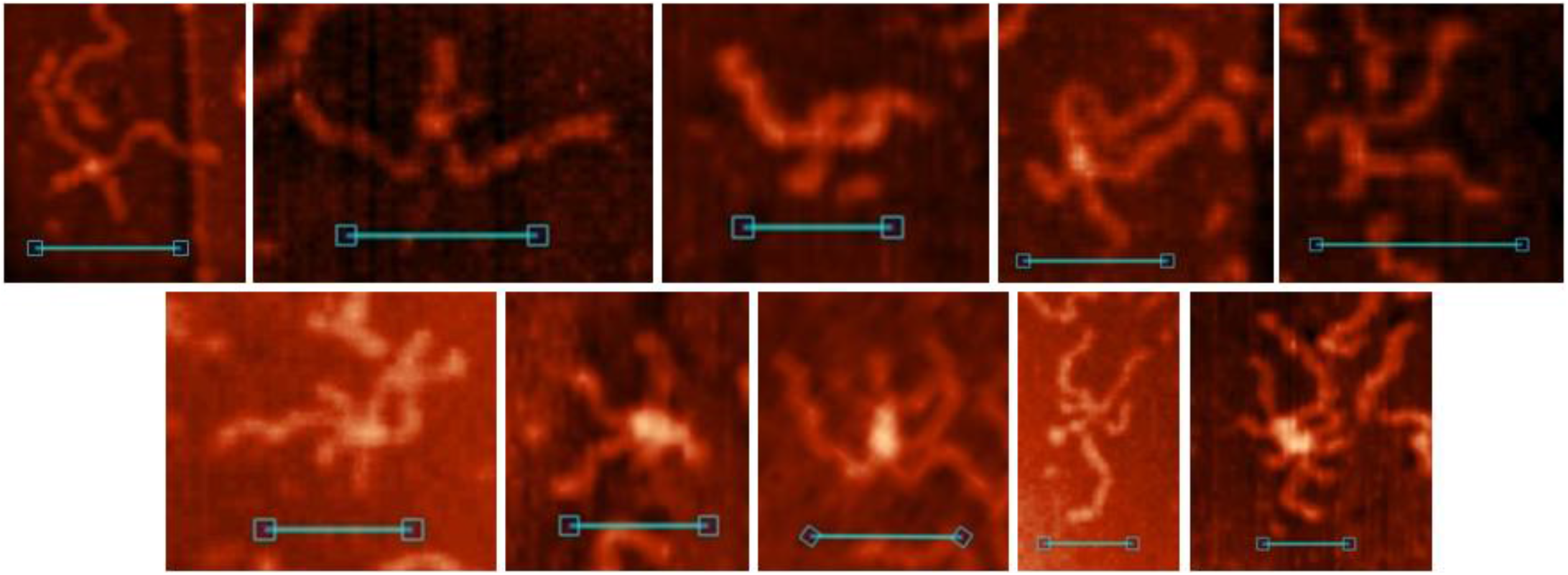
Di- and multimeric G4/IM-synaptic complexes, formed by 5s sample.

In all molecular models, G4s have a parallel conformation, because this is a conformation of the (G_3_T)_3_G_3_ quadruplex at the single-strand level (48). A molecular dynamic simulation was performed (4 ns) for all the molecular models presented here, which proved their stability.

### Antibody analysis of the G4/IM-synaptic complexes

The above G4/IM-synaptic complex structures form spontaneously from separate DNA duplexes, as determined by PCR. This method excludes Holliday structure (HS) formation, but one may note that, although our buffer does not contain Mg^2+^ ions, many of the synaptic structures are similar to HS in a stacked conformation (49) on AFM images. Antibody analysis was used to confirm the presence of G4 in the synaptic complexes.

We checked the affinity of the anti-G4 DNA antibody (clone 1H6; 24, 25) to model G4/IM-synaptic complex forming sequences (2m–6m) and synthesised immobile HS (see Materials and methods and Fig. S5). The 1H6 antibody (globes with height 3.5±0.5 nm and diameter 8–17 nm) does not interact with the extended conformation (dominates in AFM buffer, Fig. 9A) and stacked forms (realised in presence of 10 mM Mg^2+^, Fig. 9B) of Holliday junction, but recognises most folded G4/IM-synaptic complex molecules for all 2m – 6m samples. The AFM images show the interaction of the antibody with the intersection points of cruciforms (Fig. 9C) and higher-order structures (Fig. 9D). One or two antibody molecules interact with cruciforms, which confirms the formation of a synaptic structure via G4–G4 stacking and is observed for synaptic complexes formed by 4m, 5m, and 6m, but not by 2m and 3m. Not every cruciform was recognised by antibody molecules supposedly due to hiding of G4 within the synaptic complex structure or folding via IMs and absence of G4 motifs in these associates. We also observed HS formation at extended conformation (they are clearly distinguishable from synaptic complexes) in the AFM images, which has been described later.

**Fig. 9.**
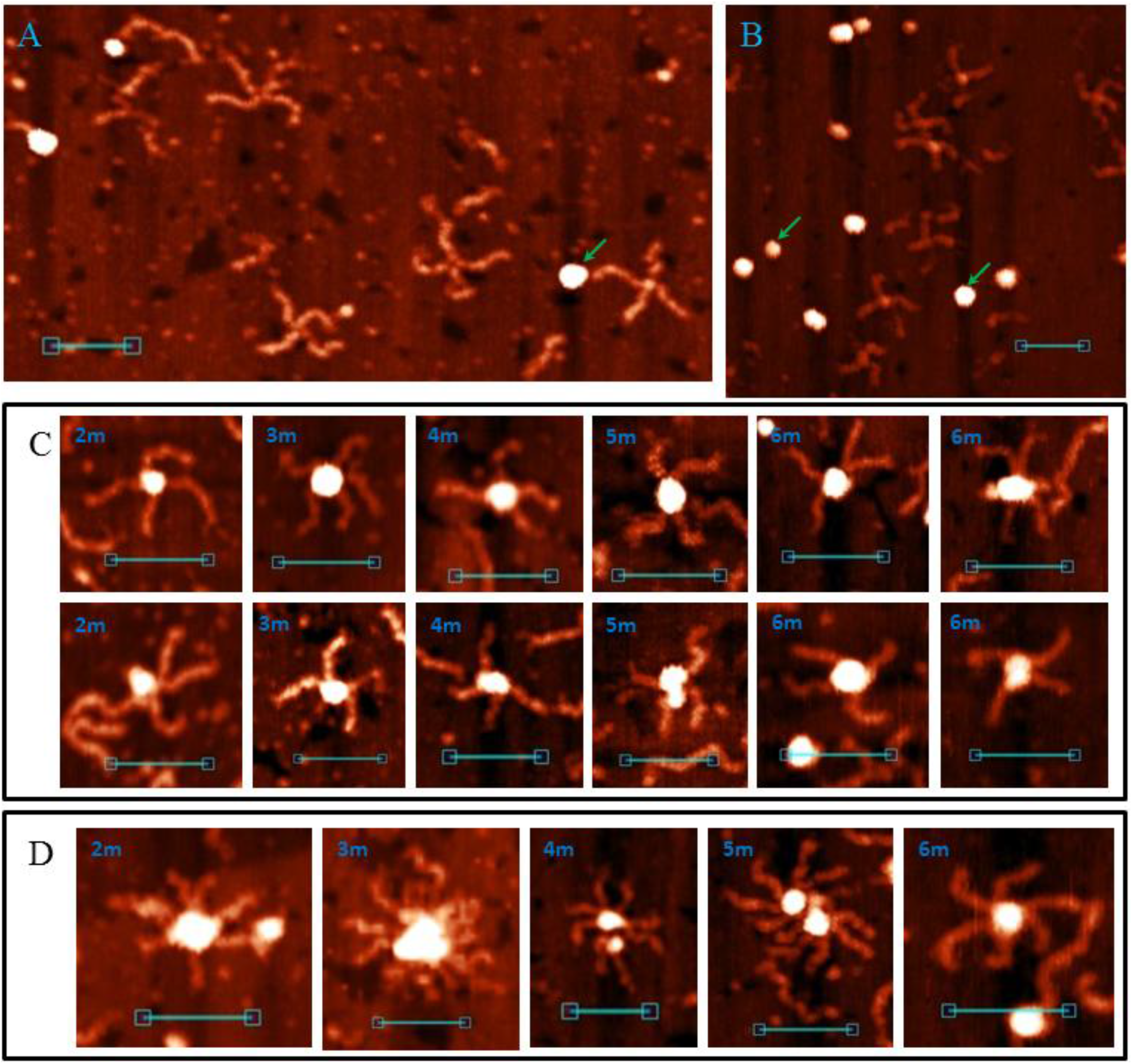
Interaction of the G4-specific antibody (1H6) with G4/IM-synaptic complexes and Holliday structure (HS). **A**. AFM image of HS in extended conformation in the presence of 1H6 antibody; **B**. AFM image of HS in a stacked form in the presence of 1H6 antibody; **C**. Interaction of 1H6 antibody with cruciform G4/IM-synaptic complexes; **D**. Interaction of 1H6 antibody with higher order synaptic complexes. Antibody molecules are marked with arrows.

We also found that the G4-DNA antibody colocalises to the central regions of some duplexes, which do not participate in G4/IM-synaptic complex formation. This is similar to the confirmation of G4 folding in dsDNA, but statistically it happens not more often than random colocalisation of antibody molecules at other parts of DNA molecules. Therefore, we may conclude that formation of the G4 structure in dsDNA is more likely when it folds as a part of the synaptic complex than by itself.

### G4/IM-synaptic complexes and recombination

As mentioned previously, G4/IM-synaptic complexes were not the only structures observed in the AFM images (Fig. 10). Cruciform junctions with lengthy intersection points, “stretched bodies” (Fig. 10A); molecular formations with two distinct HSs (Fig. 10B); and finely two DNA molecules connected by just one HS (Fig. 10C) were also observed. The height of “stretched bodies” is ~1.2 nm, and their length varies from 10 to 35 nm (shorter “stretched bodies” are not distinguishable from the IM-based synaptic complexes in the AFM images). HSs in Fig. 10B are often stacked despite the lack of Mg^2+^ ions in AFM buffer. This may be due to the steric hindrance of the two extended HSs formed quite near each other. At junctions through one HS, they are always in an extended conformation. The formation of these structures was observed for every DNA duplex containing PQS (but not for the 0 and 0Myc samples). By analysing these images, we may suppose that they show different snapshots of one process – G4/IM-synaptic complex-mediated strand exchange (*in vitro* recombination). The suggested schematic mechanism for this process is shown in Fig. 10D. The first step is the formation of base pairs (here AT pairs) between G4 or IM loops and free complimentary chains of the other duplex from the synaptic complex. Next, disrupting the synaptic complex structure leads not to the recovery of the initial two duplexes, but confusing between chains and the formation of two HSs. Their migration away from each other under *in vitro* conditions (in the absence of proteins mediating this process, such as RuvAB from *E. coli*), causes negative superhelicity of the DNA ring bordered by the HSs. Therefore, we may conclude that duplexes within HSs (Fig. 10B) are partly unwind, which contribute to the folding of noncanonical structures, for example, emergence of Z-DNA areas or synaptic complexes (as in the 6m sample, bottom image). Such folds with two distinct HSs are extremely rare and most molecules with negative superhelicity are supercoiled (Fig. 10A) with different lengths and positions of the supercoiled part, which is consistent with spontaneous branch migration. Finally, when one HS reaches the end of the duplexes and resolves, superhelicity will be lost and the duplexes will be joined by one HS in an extended conformation (Fig. 10C).

**Fig. 10.**
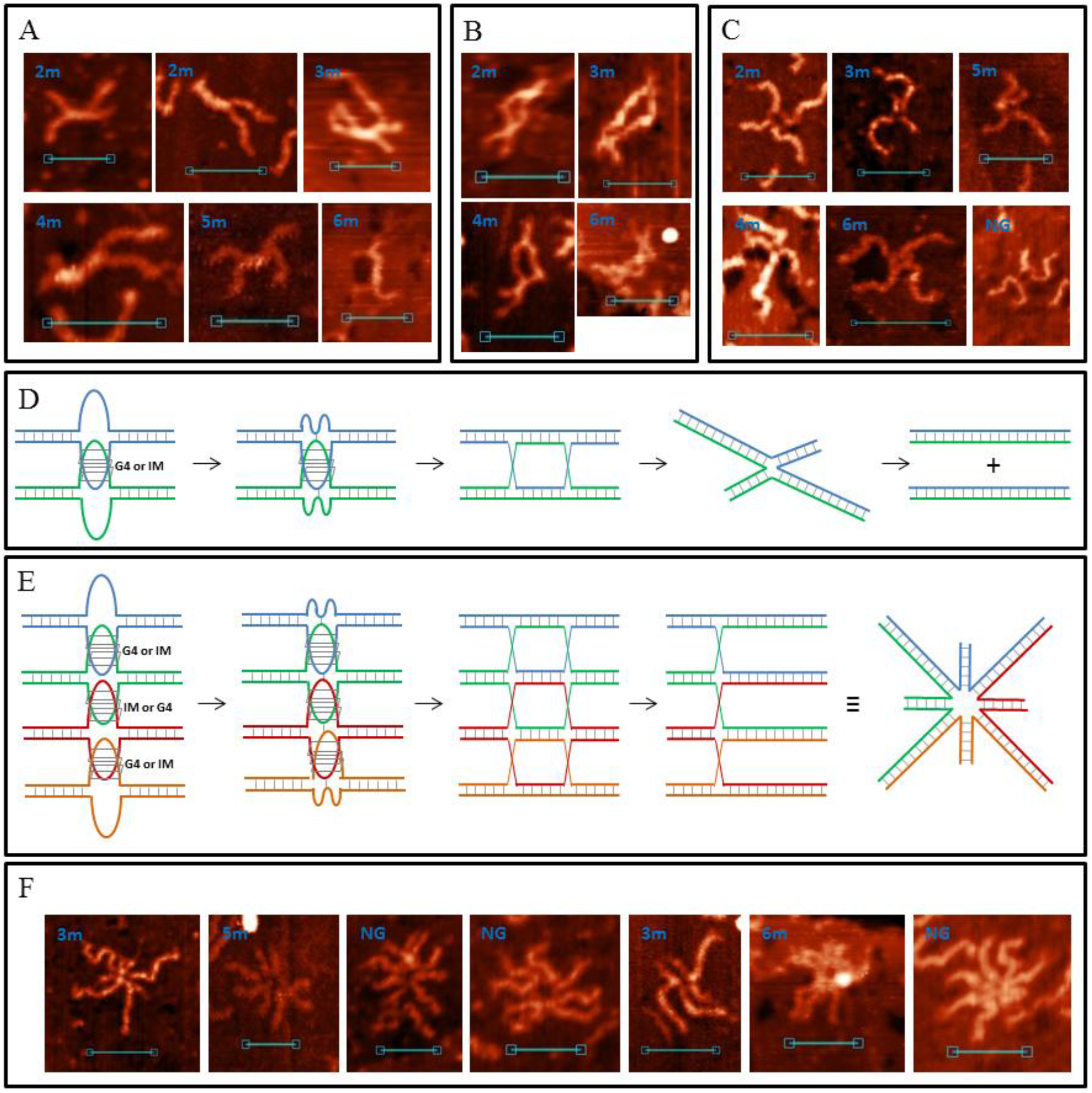
G4/IM-synaptic complex-promoted DNA strand exchange (recombination). **A**. Examples of junctions with lengthy intersection points, revealed by AFM. **B**. Structures with two HSs. **C**. Junctions through one HS. Scale bars: 50 nm. **D**. Schematic representation of the presumed mechanism of G4/IM-synaptic complex-promoted DNA strand exchange. **E**. Presumed mechanism of tetra-HS formation from tetrameric G4/IM-synaptic complex. **F**. Examples of multi-Holliday structures revealed by AFM. Scale bars: 50 nm.

Analogously, we suggest that base pair formation between converged G4 or IM loops and complementary free chains from the other duplexes may also occur at multimeric synaptic complexes. Subsequent refolding of its structure and confusion between chains must lead to the formation of a multi-HS. This process for a four-duplex synaptic complex is schematically depicted in Figure 10E. We write about this multi-recombination process and multi-HS formation because such associates form even if extremely rare. All the AFM images are shown in Fig. 10F. Most of them (three multi-HSs) are formed by NG sample, which is a 200 bp amplicon of G4-containing sequence located upstream of the *N. gonorrhoeae* pilin expression locus (pilE) and necessary for initiation of nonreciprocal recombination between pilE and one of many silent pilin loci [7]. Based on our results, we may suggest that the process of pilin antigen switching goes through multimeric synaptic complex formation and its resolution to multi-HS. In the multi-Holliday structure (as well as in usual HS) in the extended conformation, long and short duplex arms must alternate (Fig. 10E), but as clearly seen (particularly for the 3m sample), this does not always occur. Presumably, they are not fully extended but are partly stacked.

## DISCUSSION

We showed that DNA duplexes containing PQS may spontaneously form G4/IM-synaptic complexes. Such structures can be formed even by truncated PQS sequences; therefore there are more synaptic-forming sequences than predicted using different G4-finding software. Molecular modelling was used to elucidate the complex fine structures and the results confirmed the structures observed using AFM. G4 folding was confirmed through experiments with an anti-G4 DNA antibody. The mechanism of G4/IM-synaptic complex-mediated recombination was proposed.

The mechanism of synaptic complex formation should be further investigated. AFM experimental conditions – low ionic strength, particularly K^+^ concentration – do not contribute to G4 and IM folding. However, it also reduces duplex stability, which may contribute to synaptic complex folding. *In vivo* conditions (≈100 mM KCl, molecular crowding) stabilise duplexes and synaptic complexes. Synaptic complex folding requires DNA duplex melting. The formation of right-handed crossovers may trigger such melting and lead to noncanonical secondary structure folding (50, 51). Most right-handed crosses require cytosine–phosphate group interactions at the anchoring point and are frequently stabilised by divalent cations (usually Mg^2+^). Our AFM buffer did not contain Mg^2+^ ions, so we propose that this lack is compensated by cytosine multiplicity at PQS sites, but adding 10 mM MgCl_2_ to AFM buffer did not increase synaptic complex formation.

AFM spectroscopy showed that only about percentages of duplexes containing PQS partially fold into G4/IM-synaptic complexes, but we also observed examples of varying degrees of synaptic complex decay (Fig. S6). This means that their real quantity in the solution may be greater than that observed on the substrate. It is clearly visible that G/C-rich ssDNA chains are temporarily released during synaptic complex decomposition. Their formation may contribute to genome instability and DSB formation.

In this study, we correlate AFM and molecular modelling data for only one simple G4/IM-pair (with single-base identical loops and only parallel G4 conformation). The diversity of synaptic complex structures is defined by the PQS sequence. For example, diversity of possible synaptic complexes, formed by duplexes containing telomeric PQS, must increase because telomeric G4 is polymorphic (depending on the local environment, it may fold into parallel, antiparallel, or hybrid-type structures) [3]. The possibility of forming synaptic complexes by duplexes containing different PQSs (more probable cases in natural conditions), additionally diversifies possible structures.

One duplex pair may form diverse synaptic structures, and the tendency to form one or another type depends on the conditions, initial mutual arrangement, and sequences. Different folded structures may bind to different protein factors, so synaptic complex heterogeneity may be a factor of uncertainty, and also serves as a fine-tuning element of biochemical processes.

Usually complementary NA–NA interactions are studied, but this study proves that DNA molecule may be not only a passive information carrier, which is realised by protein factors, but also directly influences many regulatory processes in cell life due to DNA secondary structure polymorphism. The *in vivo* existence of G4/IM-synaptic complex awaits bona fide verification, but their folding may facilitate transient DNA–DNA contacts, contribute to shaping the long-standing chromatin organisation, and participate as the initiating stage of important processes such as homologous and non-homologous recombination, enhancer/suppressor activity, translocation, chromatin remodelling, and DSB formation.

## SUPPLEMENTARY DATA

Supplementary Data are available at NAR online.

## ACKNOWLEDGEMENT

The authors express their sincere gratitude to Dr. M.A. Lagarkova and Malakhova M.V. for providing human and *N. gonorrhoeae* total genomic DNA and A.M. Varizhuk for her comments and fruitful discussion of the material. We would like to thank Editage (www.editage.com) for English language editing.

## FUNDING

This work was supported by the Russian Foundation for Basic Research [19-015-00024]. Funding for open access charge: Russian Foundation for Basic Research.

## CONFLICT OF INTEREST

None declared.

## Notes

### Competing Interest Statement

The authors have declared no competing interest.

