## Supplemental Data 1 for "Spontaneous DNA synapsis by forming noncanonical intermolecular structures"

**Table S1**. Sequences of natural and model DNA-duplexes used in AFM experiments. Yellow highlighted are PQSs and their complimentary C-rich chains.

| | |
|---|---|
| cMyc | 5’AATGGTAGGCGCGCGTAGTTAATTCATGCGGCTCTCTTACTCTGTTTACATCCTAGAGCTAGAGTGCTCGGCTGCCCGGCTGAGTCTCCTCCCCACCTTCCCCACCCTCCCCACCCTCCCCATAAGCGCCCCTCCCGGGTTCCCAAAGCAGAGGGCGTGGGGGAAAAGAAAAAAGATCCTCTCTCGCTAATCTCCGCCCAC<br>3’TTACCATCCGCGCGCATCAATTAAGTACGCCGAGAGAATGAGACAAATGTAGGATCTCGATCTCACGAGCCGACGGGCCGACTCAGAGGAGGGGTGGAAGGGGTGGGAGGGGTGGGAGGGGTATTCGCGGGGAGGGCCCAAGGGTTTCGTCTCCCGCACCCCCTTTTCTTTTTTCTAGGAGAGAGCGATTAGAGGCGGGTG |
| kRas | 5’CGCAGAGGGCAGAGCTATCGATGCGTTCCGCGCTCGATTCTTCTTCAGACGGGCGTACGAGAGGGAGCGGCTGAGGGCGGTGTGGGAAGAGGGAAGAGGGGGAGGCAGCGAGCGCCGGCGGGGAGAAGGAGGGGGCCGGGCCGGGCCGGCGGGGGAGGAGCGGGGGCCGGGCCGGCGGAGGAAGGGGTGGCTGGGGCGGTCTAGGGTG<br>3’GCGTCTCCCGTCTCGATAGCTACGCAAGGCGCGAGCTAAGAAGAAGTCTGCCCGCATGCTCTCCCTCGCCGACTCCCGCCACACCCTTCTCCCTTCTCCCCCTCCGTCGCTCGCGGCCGCCCCTCTTCCTCCCCCGGCCCGGCCCGGCCGCCCCCTCCTCGCCCCCGGCCCGGCCGCCTCCTTCCCCACCGACCCCGCCAGATCCCAC |
| NG | 5’CAGTTTTAGTGCCGATTTTCGTTACTTTTTTATTGGCATGGGGTATCGGGTGTGTTGATTGGGTCGGAATTTGAGATTTTTGAATTTACGCGTTAGAATAGGGTGGGTTGGGTGGGGAATTTTCTATTTTTTAAAAAGCTCCGTTTTCTTGGAAAGCATTTGAAATCGGCGCGTGGTGTTTCTATGCAACCGGCAGATGA<br>3’GTCAAAATCACGGCTAAAAGCAATGAAAAAATAACCGTACCCCATAGCCCACACAACTAACCCAGCCTTAAACTCTAAAAACTTAAATGCGCAATCTTATCCCACCCAACCCACCCCTTAAAAGATAAAAAATTTTTCGAGGCAAAAGAACCTTTCGTAAACTTTAGCCGCGCACCACAAAGATACGTTGGCCGTCTACT |
| 0Myc | 5’CACGGAAGTAATACTCCTCTCCTCTTCTTTGATCAGAATCGATGCATTTTTTGTGCATGACCGCATTTCCAATAATAAAAGGGGAAAGAGGACCTGGAAAGGAATTAAACGTCCGGTTTGTCCGGGGAGGAAAGAGTTAACGGTTTTTTTCACAAGGGTCTCTGCTGACTCCCCCGGCTCGGTCCACAAGCTCTCCACTT<br>3’GTGCCTTCATTATGAGGAGAGGAGAAGAAACTAGTCTTAGCTACGTAAAAAACACGTACTGGCGTAAAGGTTATTATTTTCCCCTTTCTCCTGGACCTTTCCTTAATTTGCAGGCCAAACAGGCCCCTCCTTTCTCAATTGCCAAAAAAAGTGTTCCCAGAGACGACTGAGGGGGCCGAGCCAGGTGTTCGAGAGGTGAA |
| 0 | 5’GTAGTGAGACTCCTGAAAGAAGTATCTGACCAACTTACAGATAATATTAAAGCTCTACACGAGACCTCCAATGTGATCAGCTGCACGTATCTGACCAACTTACAGATAAGCCAAGTGTGATCAGCTGCACGTCATGTTCCTGACTTTAGACGTATCTGACCAACTTACAGATAAGGAATCCATCTAGACTCAGCG<br>3’CATCACTCTGAGGACTTTCTTCATAGACTGGTTGAATGTCTATTATAATTTCGAGATGTGCTCTGGAGGTTACACTAGTCGACGTGCATAGACTGGTTGAATGTCTATTCGGTTCACACTAGTCGACGTGCAGTACAAGGACTGAAATCTGCATAGACTGGTTGAATGTCTATTCCTTAGGTAGATCTGAGTCGC |
| 2m | 5’GTAGTGAGACTCCTGAAAGAAGTATCTGACCAACTTACAGATAATATTAAAGCTCTACACGAGACCTCCAATGTGATCAGCTGCACGTATCTGACCCACCCACAGATAAGCCAAGTGTGATCAGCTGCACGTCATGTTCCTGACTTTAGACGTATCTGACCAACTTACAGATAAGGAATCCATCTAGACTCAGCG<br>3’CATCACTCTGAGGACTTTCTTCATAGACTGGTTGAATGTCTATTATAATTTCGAGATGTGCTCTGGAGGTTACACTAGTCGACGTGCATAGACTGGGTGGGTGTCTATTCGGTTCACACTAGTCGACGTGCAGTACAAGGACTGAAATCTGCATAGACTGGTTGAATGTCTATTCCTTAGGTAGATCTGAGTCGC |
| 3m | 5’GTAGTGAGACTCCTGAAAGAAGTATCTGACCAACTTACAGATAATATTAAAGCTCTACACGAGACCTCCAATGTGATCAGCTGCACGTATCTGACCCACCCACCCATAAGCCAAGTGTGATCAGCTGCACGTCATGTTCCTGACTTTAGACGTATCTGACCAACTTACAGATAAGGAATCCATCTAGACTCAGCG<br>3’CATCACTCTGAGGACTTTCTTCATAGACTGGTTGAATGTCTATTATAATTTCGAGATGTGCTCTGGAGGTTACACTAGTCGACGTGCATAGACTGGGTGGGTGGGTATTCGGTTCACACTAGTCGACGTGCAGTACAAGGACTGAAATCTGCATAGACTGGTTGAATGTCTATTCCTTAGGTAGATCTGAGTCGC |
| 4m | 5’GTAGTGAGACTCCTGAAAGAAGTATCTGACCAACTTACAGATAATATTAAAGCTCTACACGAGACCTCCAATGTGATCAGCTGCACGTATCCCACCCACCCACCCATAAGCCAAGTGTGATCAGCTGCACGTCATGTTCCTGACTTTAGACGTATCTGACCAACTTACAGATAAGGAATCCATCTAGACTCAGCG<br>3’CATCACTCTGAGGACTTTCTTCATAGACTGGTTGAATGTCTATTATAATTTCGAGATGTGCTCTGGAGGTTACACTAGTCGACGTGCATAGGGTGGGTGGGTGGGTATTCGGTTCACACTAGTCGACGTGCAGTACAAGGACTGAAATCTGCATAGACTGGTTGAATGTCTATTCCTTAGGTAGATCTGAGTCGC |
| 5m | 5’GTAGTGAGACTCCTGAAAGAAGTATCTGACCAACTTACAGATAATATTAAAGCTCTACACGAGACCTCCAATGTGATCAGCTGCACGTCCCACCCACCCACCCACCCAAGCCAAGTGTGATCAGCTGCACGTCATGTTCCTGACTTTAGACGTATCTGACCAACTTACAGATAAGGAATCCATCTAGACTCAGCG<br>3’CATCACTCTGAGGACTTTCTTCATAGACTGGTTGAATGTCTATTATAATTTCGAGATGTGCTCTGGAGGTTACACTAGTCGACGTGCAGGGTGGGTGGGTGGGTGGGTTCGGTTCACACTAGTCGACGTGCAGTACAAGGACTGAAATCTGCATAGACTGGTTGAATGTCTATTCCTTAGGTAGATCTGAGTCGC |
| 6m | 5’GTAGTGAGACTCCTGAAAGAAGTATCTGACCAACTTACAGATAATATTAAAGCTCTACACGAGACCTCCAATGTGATCAGCTGCACCCCACCCACCCACCCACCCACCCGCCAAGTGTGATCAGCTGCACGTCATGTTCCTGACTTTAGACGTATCTGACCAACTTACAGATAAGGAATCCATCTAGACTCAGCG<br>3’CATCACTCTGAGGACTTTCTTCATAGACTGGTTGAATGTCTATTATAATTTCGAGATGTGCTCTGGAGGTTACACTAGTCGACGTGGGGTGGGTGGGTGGGTGGGTGGGCGGTTCACACTAGTCGACGTGCAGTACAAGGACTGAAATCTGCATAGACTGGTTGAATGTCTATTCCTTAGGTAGATCTGAGTCGC |
| 5s | 5’GTAGTGAGACTCCTGAAAGAAGTCCCACCCACCCACCCACCCAATATTAAAGCTCTACACGAGACCTCCAATGTGATCAGCTGCACGTATCTGACCAACTTACAGATAAGCCAAGTGTGATCAGCTGCACGTCATGTTCCTGACTTTAGACGTATCTGACCAACTTACAGATAAGGAATCCATCTAGACTCAGCG<br>3’CATCACTCTGAGGACTTTCTTCAGGGTGGGTGGGTGGGTGGGTTATAATTTCGAGATGTGCTCTGGAGGTTACACTAGTCGACGTGCATAGACTGGTTGAATGTCTATTCGGTTCACACTAGTCGACGTGCAGTACAAGGACTGAAATCTGCATAGACTGGTTGAATGTCTATTCCTTAGGTAGATCTGAGTCGC |

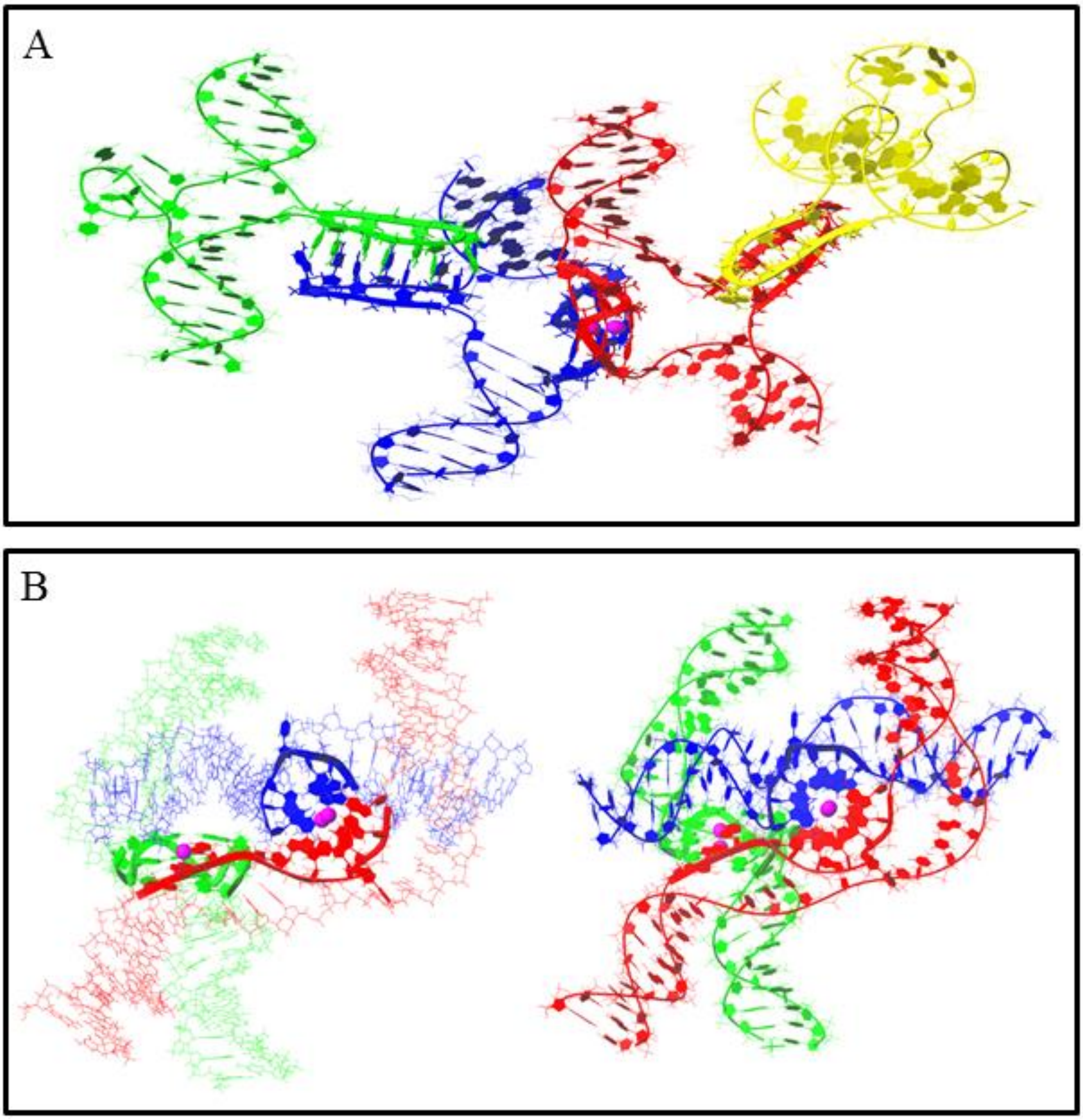


**Fig. S1. A.** Molecular model of tetrameric complex formed through intermolecular G4 and two IMs folding (not revealed by AFM). **B.** Molecular model of trimolecular synaptic complex assembled through the formation of two intermolecular G4s. Theoretically, 3m sample is able to form this structure, but it was not revealed by AFM.

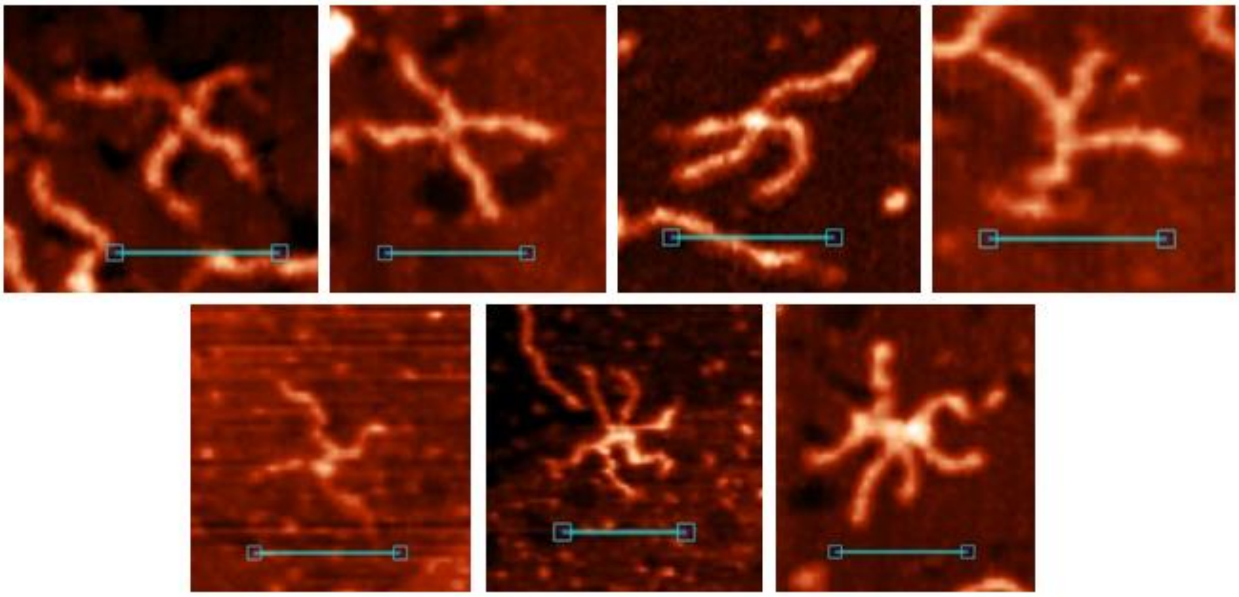

**Fig. S2.** Examples of di- and trimeric G4/IM-synaptic complexes, formed by 4m sample, with the same structures as 2m and 3m form. Scale bars: 50 nm.

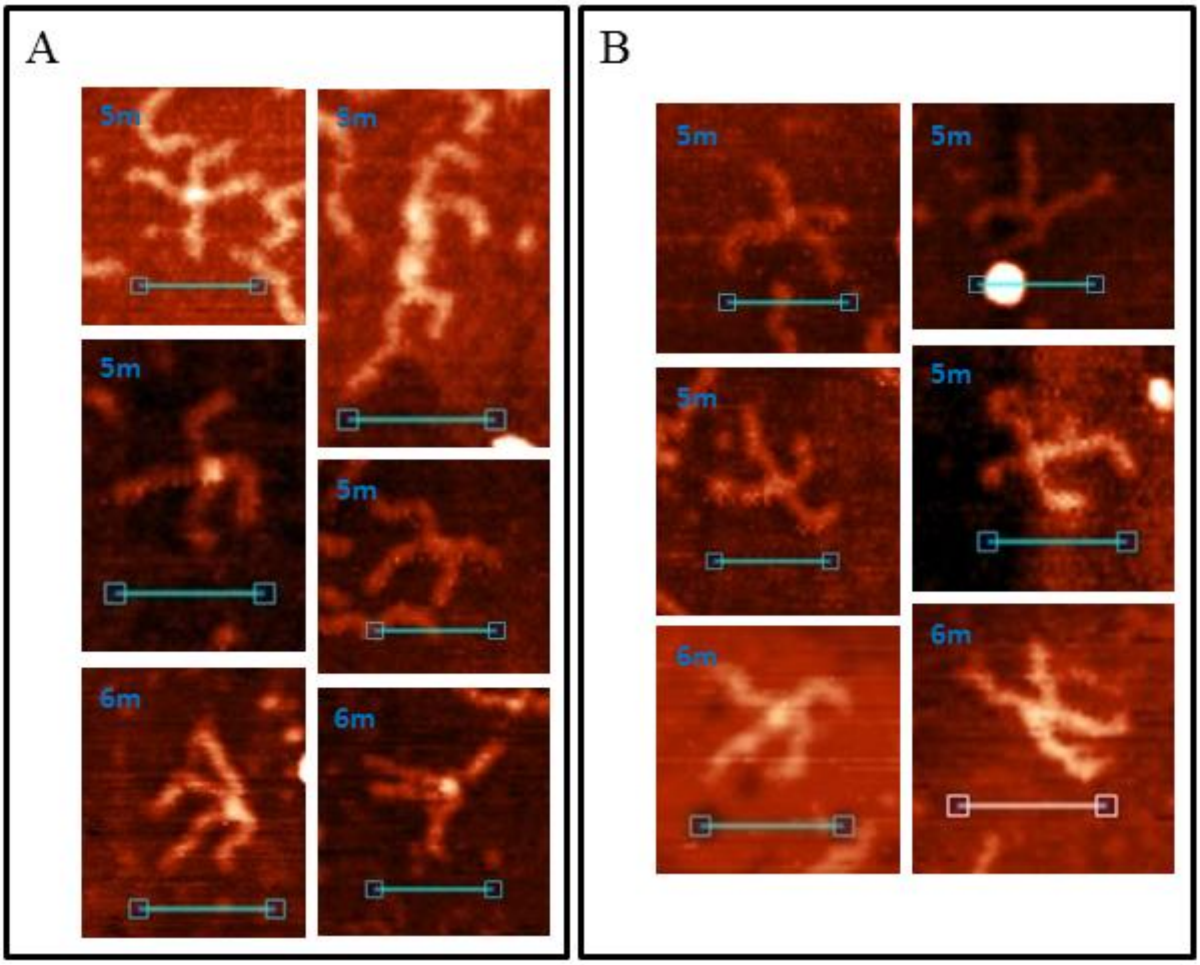


**Fig. S3.** Examples of simple cruciform synaptic complexes, formed by 5m and 6m samples. **A.** Complexes with four $G_3T$ and/or $C_3A$ blocks involved. They have the same structures as the complexes formed by 4m sample. **B.** Complexes with only two $G_3T$ and/or $C_3A$ blocks involved. They have the same structures as formed by 2m and 3m samples. Scale bars: 50 nm.

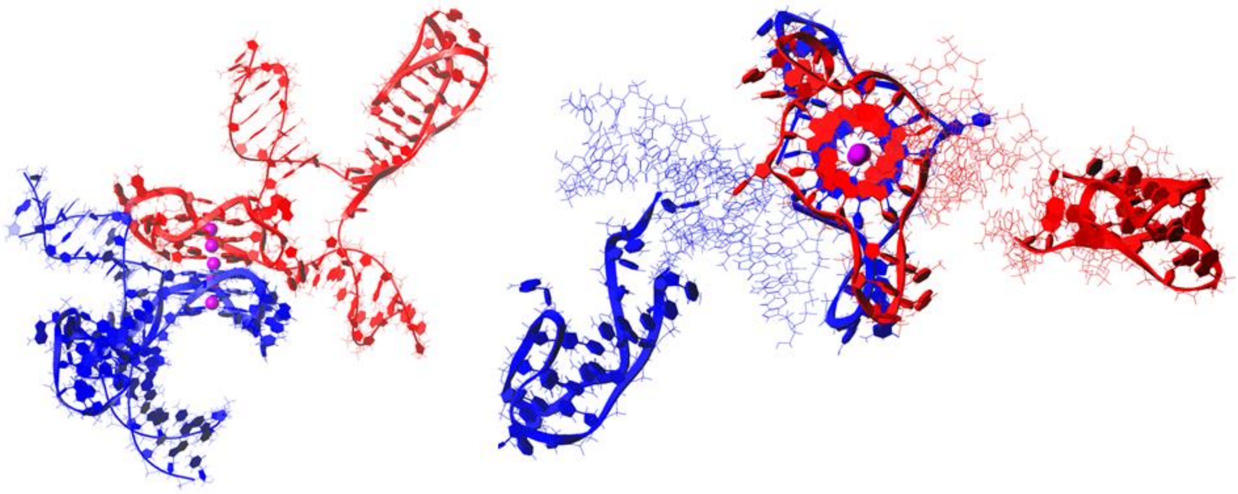

**Fig. S4.** Molecular model of the possible complex, formed by 6m sample through stacking of intramolecular G4s and containing two 5-base loops at each chain (not revealed by AFM).

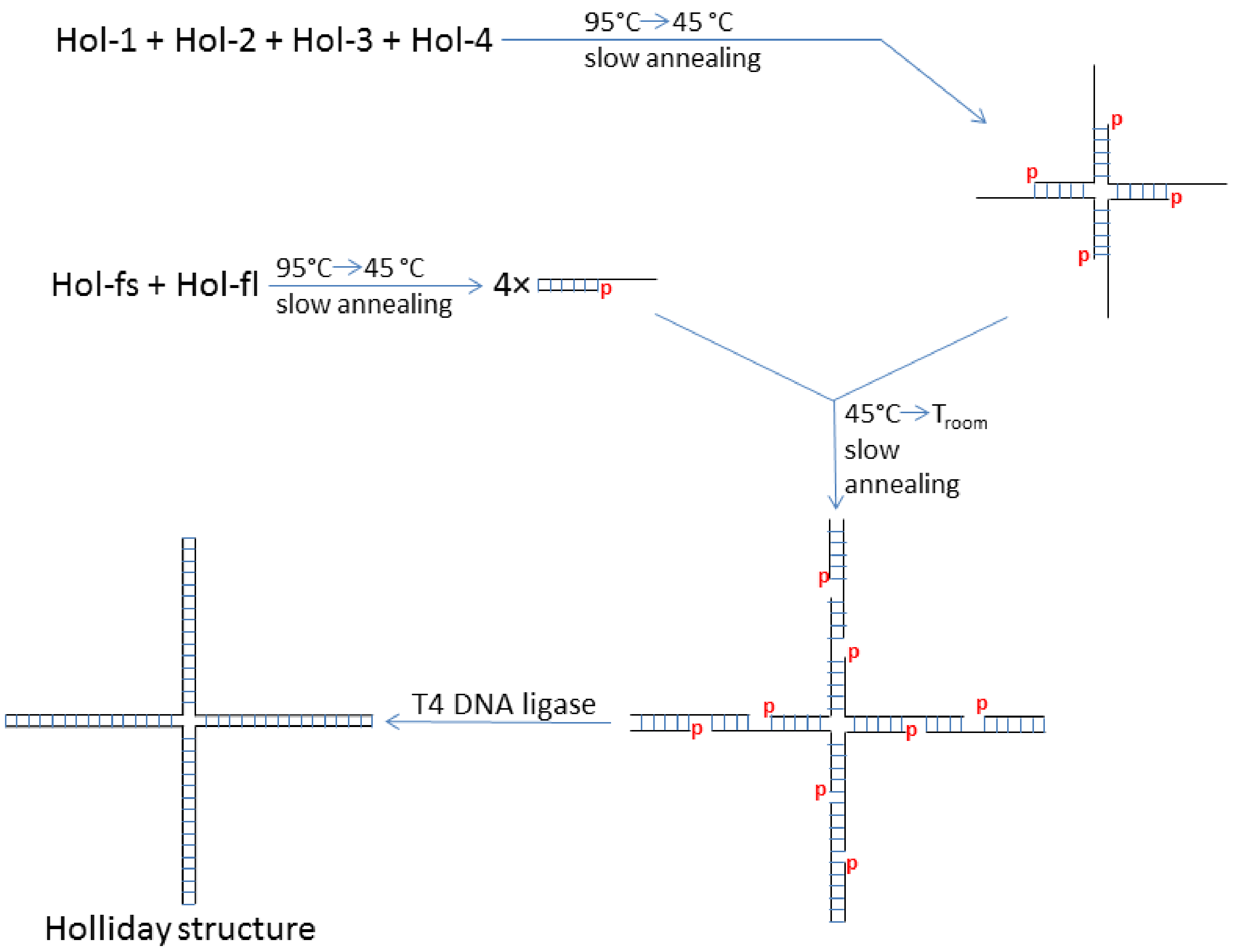


**Fig. S5.** Scheme of immobile Holliday structure synthesis.

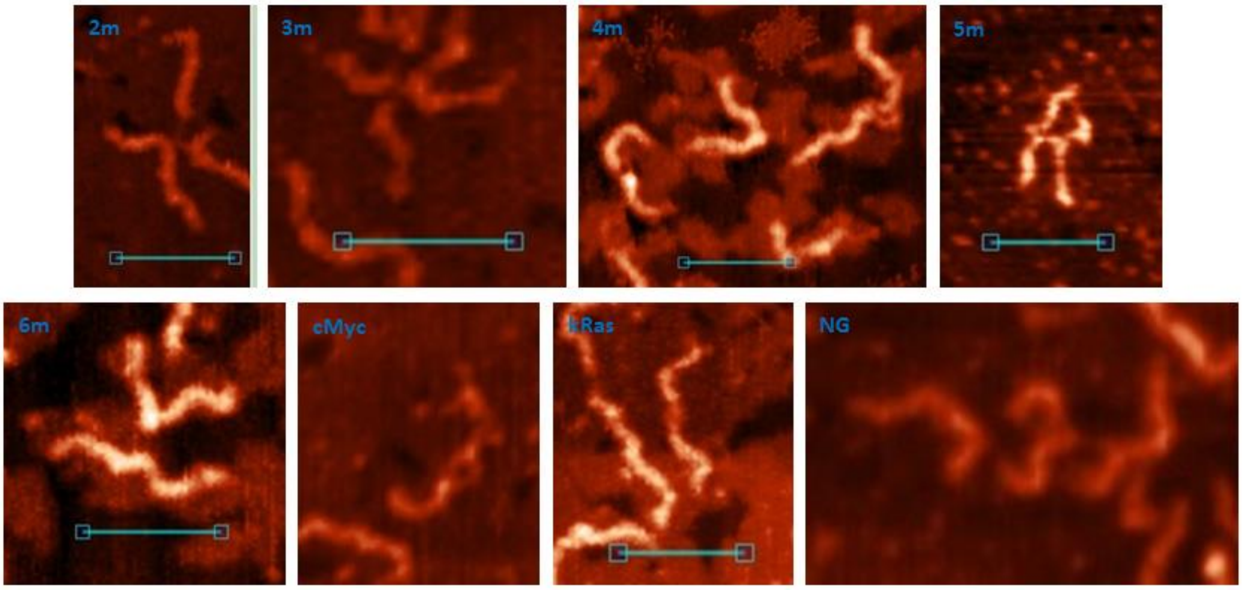


**Fig. S6.** Examples of broken G4/IM-synaptic complexes.
